# Emergence of multiple variants of SARS-CoV-2 with signature structural changes

**DOI:** 10.1101/2020.04.26.062471

**Authors:** Debaleena Bhowmik, Sourav Pal, Abhishake Lahiri, Arindam Talukdar, Sandip Paul

## Abstract

This study explores the divergence pattern of SARS-CoV-2 using whole genome sequences of the isolates from various COVID-19 affected countries. The phylogenomic analysis indicates the presence of at least four distinct groups of the SARS-CoV-2 genomes. The emergent groups have been found to be associated with signature structural changes in specific proteins. Also, this study reveals the differential levels of divergence patterns for the protein coding regions. Moreover, we have predicted the impact of structural changes on a couple of important viral proteins via structural modelling techniques. This study further advocates for more viral genetic studies with associated clinical outcomes and hosts’ response for better understanding of SARS-CoV-2 pathogenesis enabling better mitigation of this pandemic situation.

## Introduction

The recent pandemic due to novel coronavirus disease (COVID-19) originated from Wuhan, Hubei province of China has affected more than 200 countries worldwide. On December 31^st^ 2019 China reported to WHO about the infection and on January 7^th^ the causative agent was identified as the novel coronavirus or Severe Acute Respiratory Syndrome - Coronavirus 2 (SARS-CoV-2) (Surveillances V; COVID-19 situation reports). On March 11^th^ 2020, WHO declared COVID-19 as a global pandemic (Cucinotta & Vanelli, 2020). In India, the first incidence of COVID-19 was reported from the Thrissur district of Kerala (Timeline of the 2020 coronavirus pandemic in India). Apart from China, many other countries from Asia, Europe and North America are severely affected and as per the WHO situation report as of 25^th^ April there has been 2,832,454 positive cases and 198,532 deaths worldwide (COVID-19 situation reports).

SARS-CoV-2 like SARS-CoV and MERS-CoV belongs to the genus *Betacoronavirus* and has crossed the species barrier to infect humans (Letko et al., 2020; Ye et al., 2020). This single-stranded, positive-sense RNA virus shares about 79% and 50% genomic identity with that of the SARS-CoV and MERS-CoV respectively (Lu et al., 2020). Although, it was reported that SARS-CoV-2 infection may be less severe as compared to that of SARS-CoV or MERS-CoV, it is now apparent that the current novel coronavirus has an accelerated rate of human-to-human transmission than its predecessors (Wang et al., 2020; Huang et al., 2020). SARS-CoV-2 genome is around 29,903 nucleotides and organized in the following order from 5’ to 3’: ORF1ab (replicase), structural spike glycoprotein (S), ORF3a protein, structural envelope protein (E), structural membrane glycoprotein (M), ORF6 protein, ORF7a protein, ORF7b protein, ORF8 protein, structural nucleocapsid phosphoprotein (N) and ORF10 protein. ORF1ab is a large polyprotein (∼21,291 nucleotides) encoding sixteen non-structural proteins: leader protein, nsp2, nsp3, nsp4, 3C-like proteinase, nsp6, nsp7, nsp8, nsp9, nsp10, RNA-dependent RNA polymerase (RdRp), helicase, 3’-to 5’ exonuclease, endoRNAse, 2’-o-ribose methyltransferase, and nsp11 (Wu et al., 2020). Studies have revealed that the highly mutable spike (S) protein of the virus is associated with the elevated human-to-human transmission rate through interaction with ACE2 receptor of the host. S protein is one of the well-characterized proteins of the *Coronaviridae* family; this ∼1255 amino acid transmembrane protein helps the virus to attach and enter the host. Thus analysing the structure and function of this spike protein would not only unveil the pathogenicity of the virus but also might illuminate its phylogenetic origin and diversity (Babcock et al., 2004; Zhang et al., 2006; Yan et al., 2020).

Very little is known about the divergence of the SARS-CoV-2 genome. Although the earlier studies with similar RNA pathogens (e.g. HIV-1, Influenza viruses, SARS-CoV and hepatitis C virus) pointed towards the rapidly evolving nature of the viral genomes and are subjected to the strongest evolutionary forces. The ability to undergo host-specific adaptive changes in the viral genomes has enabled them to escape the innate as well as adaptive immune responses, acquire resistance towards drugs or infect new hosts (Frost et al., 2018). The rapid spread of SARS-CoV-2 around the world has enabled exposure of the virus to individuals with diverse genetic and immunological backgrounds having varied demographics (age, sex, environmental conditions, etc.) and thus potentially imposing significant selective pressures on the SARS-CoV-2 genomes. Therefore, a study highlighting the genome divergence of SARS-CoV-2 would have tremendous importance during this pandemic, providing better understanding of the genetic and phenotypic characteristics of SARS-CoV-2 pathogenesis and valuable information towards the development of drugs and vaccines.

A staggering number of genome sequences of SARS-CoV-2 are being submitted from around the world every day, and with the increasing number of positive COVID-19 cases in India, we also intended to do a comparative genomic analysis of the Indian SARS-CoV-2 isolates with respect to that of other countries. We performed the phylogenetic and other evolutionary analyses of the SARS-CoV-2 genomes, in order to reveal their shared ancestry, determine genetic diversity and detect structural mutations. This enabled us to study the divergence of this viral genome during the early phases of pandemic and identify multiple viral variants. Through computational analysis we predicted the impact of structural changes on spike glycoprotein (S). Additionally, we also studied the impact of the mutations on RdRp as it is considered to be one of the most promising target for drug development with remdesivir (Warren et al., 2016; Shannon et al., 2020), a nucleotide analog targeting RdRp. We further discussed the potential outcomes of our findings in terms of public health response in India to better control and prevent the disease.

## Methods

### Sequence collection

We considered 109 genome sequences of SARS-CoV-2 isolates from different countries. The main source of genomes is the EpiCoV database of the GISAID (www.gisaid.org) through valid registration. We collected all the 33 genomes available from India (as on April 15^th^ 2020), and the rest were obtained from among the earliest submissions from the respective countries of Europe, Asia and America (details of the country-wise genome selection provided in Table1 of the supplementary information). This working set of 109 sequences were checked for the number of ambiguous bases (N<50) and the presence of gaps (gaps<10). Also the index genome of SARS-CoV-2 (NC_045512.2) was considered from NCBI.

### Phylogenetic reconstruction and average nucleotide identity calculation

The genome alignment was performed using MUSCLE aligner in default mode present in the UGENE v34 (Edgar RC., 2004; Okonechnikov et al., 2012). The phylogenetic tree was reconstructed by using the Maximum Likelihood method and the General Time Reversible model. The initial tree for the heuristic search was obtained automatically by applying BioNJ algorithms (Tavaré & Miura., 1986; Gascuel O., 1997). Evolutionary analysis was performed using PhyML integrated in Seaview 5.0.2 (Guindon et al., 2005; Gouy et al., 2010). Further, the tree was visualized and edited with FigTree 1.4.4 (Rambaut., 2012).

The orthologous average nucleotide identity between genome sequences (OrthoANI) was measured with the help of OAU command-line tool using the USEARCH algorithm (Lee et al., 2016; Edgar RC., 2010). The resulting OrthoANI values were represented by a clustered-heat map using the online ClusVis tool (Metsalu et al., 2015).

### Genetic diversity and mutational profile analysis

In order to extract all the annotated genes from all the SARS-CoV-2 genome sequences, the isolate obtained from Wuhan, China (NC_045512.2) was used as the reference genome in our analysis. We used the standalone version of NCBI BLAST with an in-house developed perl script to fetch all the genes from each of the 109 genomes.

Using DnaSP (Rozas et al., 2017), we calculated nucleotide diversity, synonymous and non-synonymous substitution rates for each gene. Further, the TimeZone software was employed for phylogenetic analysis of each gene and detection of structural mutations in them (Chattopadhyay et al. 2013).

### Structural analysis of spike (S) glycoprotein

We performed a sequential analysis of S protein of the SARS-CoV-2 from China and India. Comparison of sequential alignment between Wuhan isolated spike sequences with 33 sequences of Indian origin was performed by Multiple Sequence Alignment (MSA) tool using Clustal Omega. Secondary structure prediction of the aligned sequences was carried out on the CFSSP (Chou and Fasman secondary structure prediction) server.

*In silico* mutation on the SARS-CoV-2 associated S protein was done on the closed state conformation (PDB ID: 6VXX). The S protein is present in a homotrimeric conditions with individual chains containing some basic loop. That loop building and modifications were done on the Swiss-Model server with respect to the Wuhan isolated spike sequence followed by analysis of these mutations by Mutprep server. The mutation on the identified protein residues was analyzed on Pymol with a pre-existing suitable rotamer library. Structural modification and kinetics change upon mutation were performed on Dynemut Server. The computational analysis figures were generated in Pymol.

## Results and Discussion

### Phylogenomic study of SARS-CoV-2 reveals the appearance of distinct phylogenetic groups

Phylogenetic reconstruction of the divergence of all 109 SARS-CoV-2 genomes isolated from different parts of the world including 33 from India is shown in Figure 1. The phylogeny indicates the emergence of new variants in this phase of the pandemic and two distinct groups of isolates emerge (Figure 1). The group A is more diverse and comprises of several sub-groups, it also includes the ancestral sequence (NC_045512.2) isolated from the patient in Wuhan, China, reported to WHO on 31/12/2019. The majority of the members in this group are from different states of China and other Asian countries although few European and USA isolates are also present. Out of 33 genomes sequenced from India, 15 sequences belong to this diverse group. The diverse nature of this group points towards the emergence of multiple viral variants within China before the spread of the virus throughout the world. The presence of genomes from other countries indicates the direct transmission of this virus from China. A detailed observation revealed the sub-group nature of this large group, implying the emergence of new variants. On the other hand, group B is more compact and comprises of European isolates and 18 Indian isolates. It is therefore quite clear that the European genomes diverged profoundly from the initial viral genomes from China and were transmitted to India. These particular isolates represent that the viral variants emerged after the start of the pandemic.

**Figure 1.**
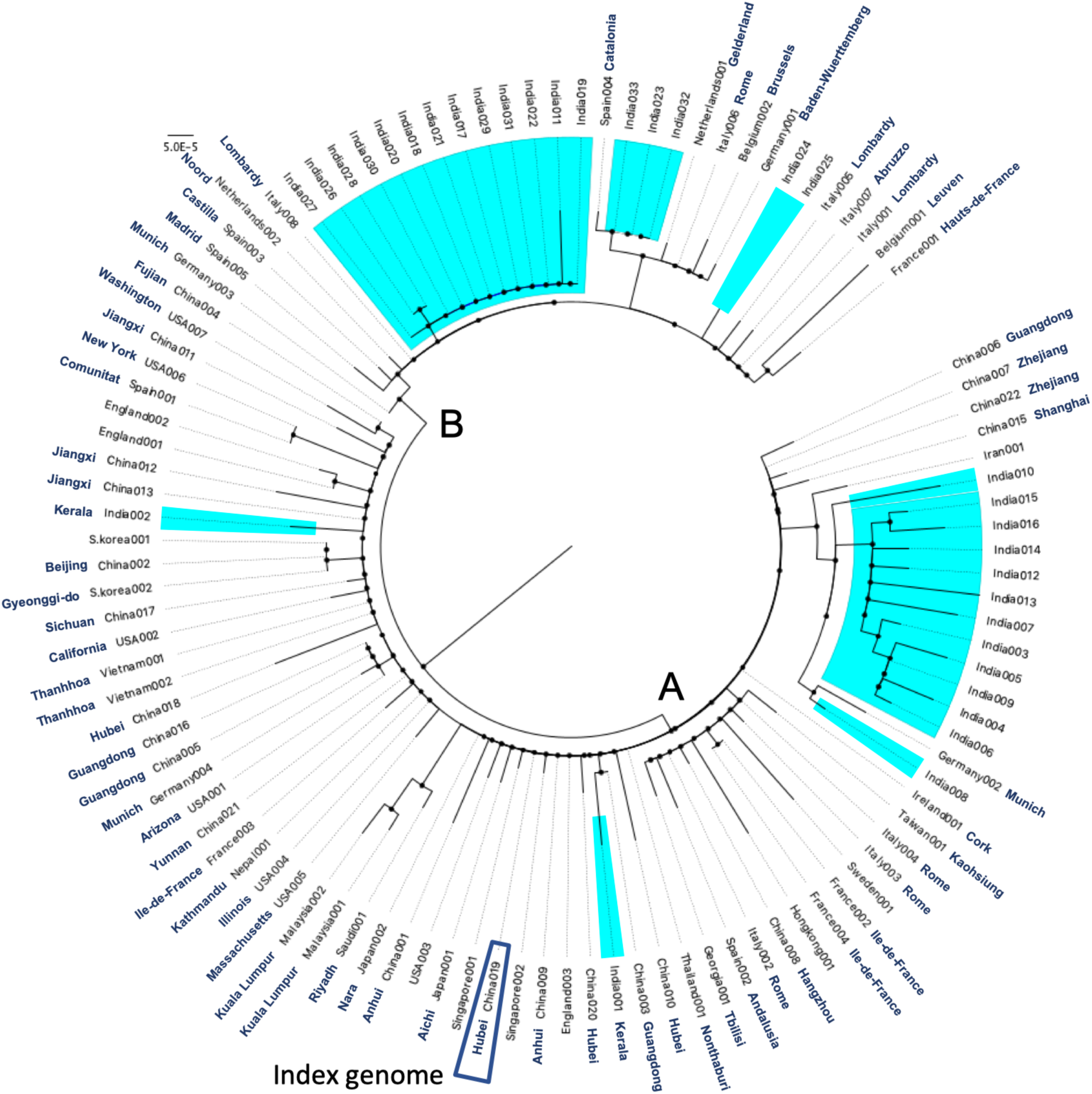
Phylogeny of 109 SARS-CoV-2 genomes. The whole genome phylogenetic tree was reconstructed by using the Maximum Likelihood method and the General Time Reversible model and tree was visualized and edited with FigTree 1.4.4. Isolates from India are marked by blue background. Available state/provinces are also depicted after the isolate names.

We further evaluated the OrthoANI values for all vs all pairs of genomes and found that the value was always more than 99.8% suggesting very close similarity. The correlation clustering of these values also indicates the segregation of isolates into two groups (A & B) and further clear sub-grouping (C, D & E) of the major group A (Figure 2). The sub-group C contains the index case of SARS-CoV-2 with other isolates from China and other countries and seems to be the ancient group that emerged in the first phase of the pandemic. This group also contains the first Indian genome sequence from Kerala. The sub-group D comprises of many Chinese isolates, European states and second Indian genome, perhaps consolidating the hypothesis stating the emergence of multiple viral variants in China before the commencement of the pandemic. Most interestingly, the sub-group E, mostly comprises of Indian isolates with genomes from Iran and Germany, probably depicting the route of transmission of viral infection in India.

**Figure 2.**
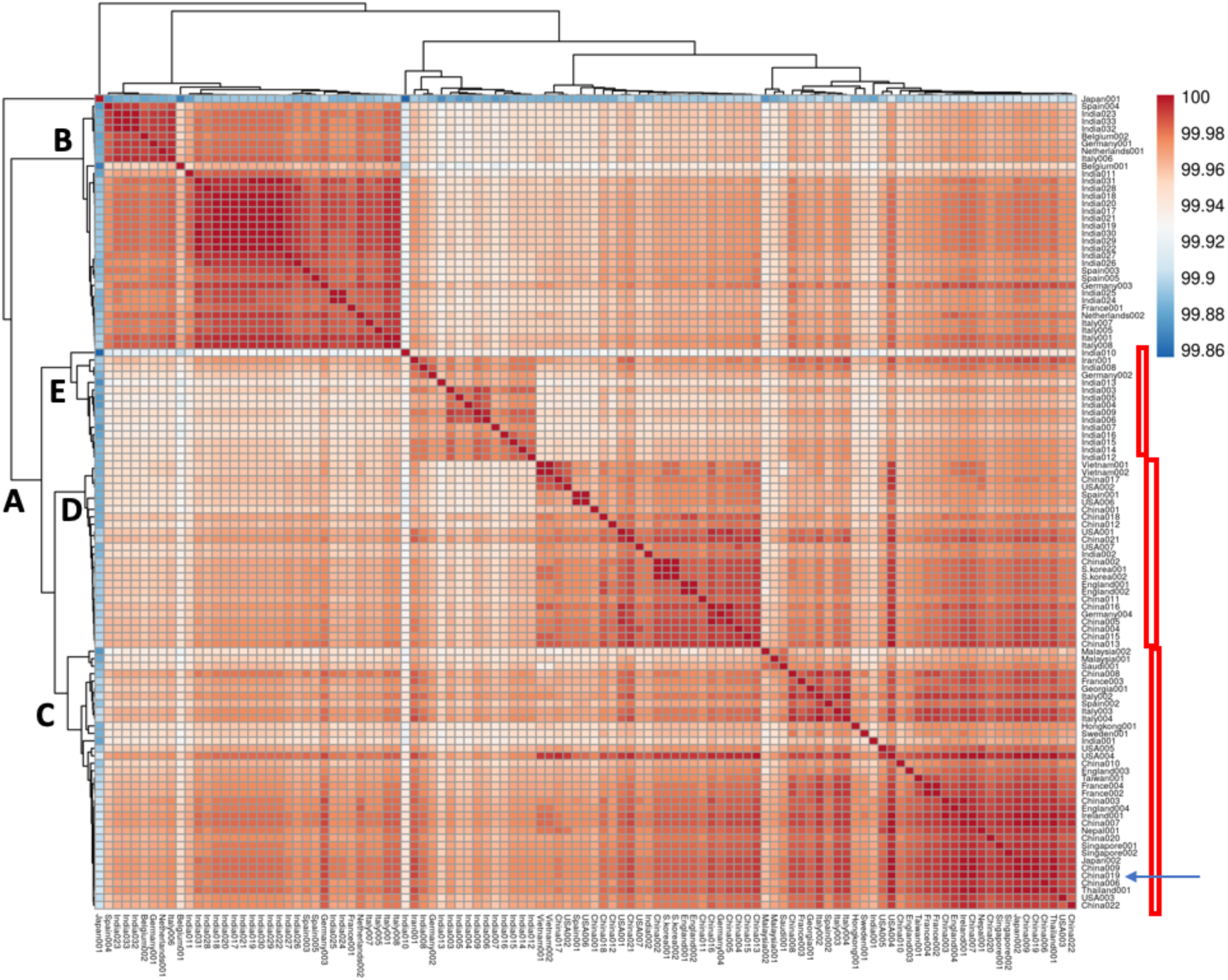
Clustering analysis of 109 genomes of SARS-CoV-2 based on the OrthoANI matrix. Both rows and columns are clustered using correlation distance and complete linkage.

The whole genome based phylogenomic analysis and subsequent nucleotide identity analysis pointed towards the presence of four distinct groups of SARS-CoV-2 genomes. It is to be mentioned here that these genomes are highly conserved as evident from >99.8% of ANI values for any pair of genomes. Coronaviruses are known to have accurate nsp14 mediated synthesis of their RNA genome and hence may maintain the overall conservation of their genomes although the nsp12 (RdRp) is not accurate (Ferron et al. 2018). The presence of multiple variants within China indicates early divergence events in early phase of the pandemic. Similar type of observations was also reported in a recent work on SARS-CoV-2 genomes (Velazquez-Salinas et al., 2020).

We also considered the available date of collection of the isolates from the metadata of the repository and observed that in sub-group C multiple early stage isolates from China are present staring from December 30^th^, 2019 collected from Wuhan, Hubei province of China. Along with, other early stage isolates from different countries are also present in this group. The first reported genome sequence from Kerala, India is also present in this group. In case of sub-group D, we again found multiple isolates from China and other countries collected between January 2020 and March 2020. The second reported genome from Kerala, India is one of the members of this particular group. Some of these isolates in different countries may emerge from the ancestors of the viral isolates although the presence of multiple viral isolates collected in the second or third week of January from China indicates the emergence of multiple variants in China itself before the initiation of the state of pandemic.

The group B isolates were collected within the month of January to March and no isolates from China can be located. These are the emergent isolates mostly originating in Europe and different from the ancestral strains and were transmitted to India. The group E is also devoid of any viral isolates from China and were collected in the month of March, 2020. These are another variant of viral isolates transmitted to India originating in Iran/Germany.

### Gene-wise genetic diversity varies between the groups

After the extraction of all the protein-coding regions from 109 genomes considering index genome (NC_045512.2) as a reference, we calculated the overall average pairwise nucleotide diversity (π) for each set. While the regions nsp8, nsp11 and ORF7b have a single variant, the regions, nsp2, nsp3, RdRp, geneS and geneN are found to have more than 10 variants. The π value for multi-variant regions varies from 4.7 × 10^−4^ to 8.8 × 10^−3^. We further compared the nucleotide diversity of Indian isolates with that of other isolates (Figure 3A). The comparison yielded several India genome-specific changes, for example, nsp15 and ORF7a showed mutational changes only for Indian isolates. On the other hand, mutations happened for nsp7, nsp9, nsp10, nsp16, geneE and ORF10 genes for some of the isolates from different parts of the world but no changes can be seen in Indian isolates.

**Figure 3.**
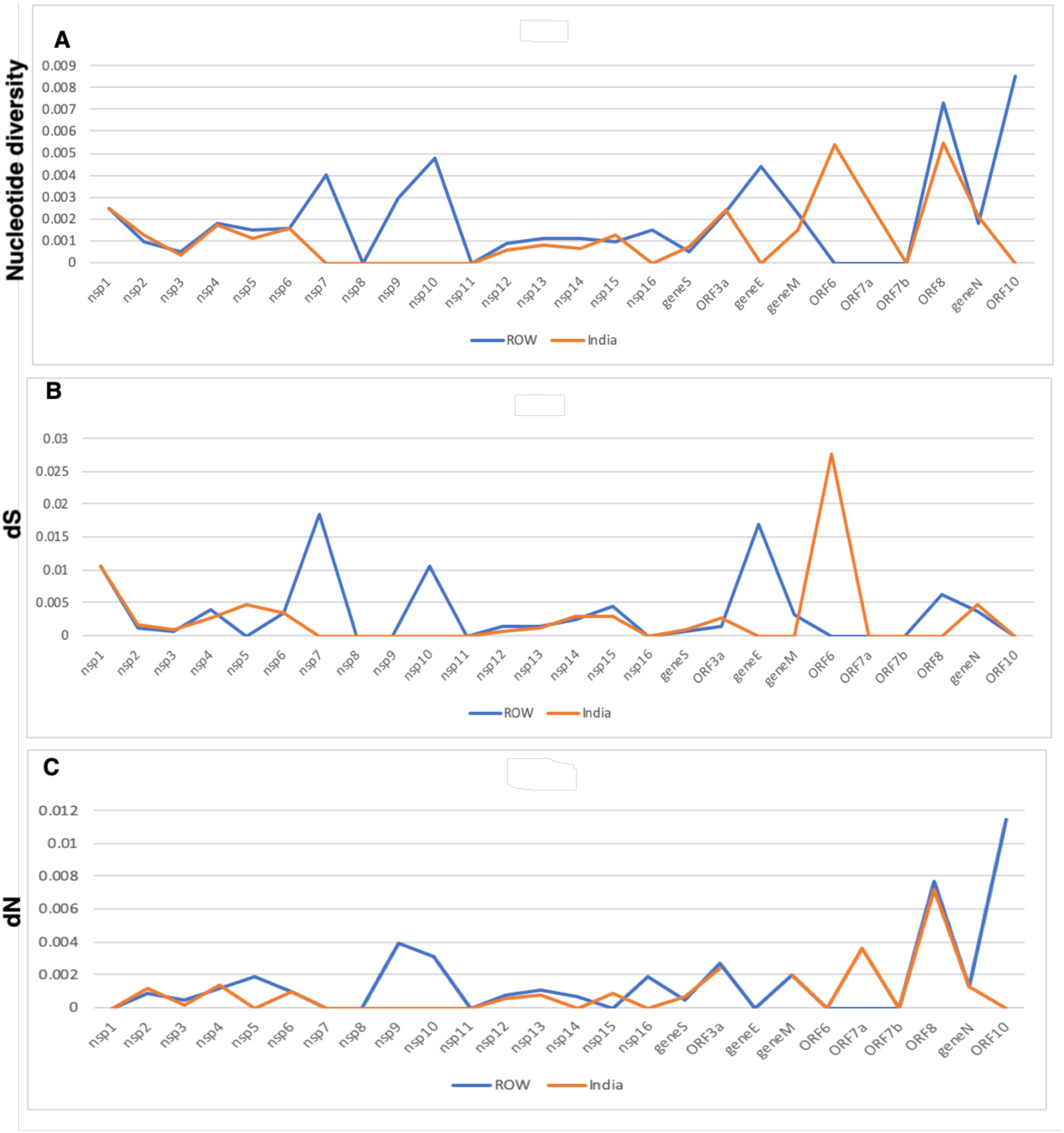
Distribution of average nucleotide diversity (A), rate of synonymous substitution per synonymous sites (B) and rate of non-synonymous substitution per non-synonymous sites for all protein-coding regions of Indian isolates (orange line) and other genomes (blue line).

Next, we estimated the rates of silent or synonymous (dS) and amino-acid changing or non-synonymous sites (dN) mutations for these sets of protein-coding regions in order to determine the types of mutations that occurred in each gene as well as if any selection pressure acting on them. The dS and dN profiles of the protein-coding regions depict selective mutational pressure for several of those regions (Figure 3B and C). The rate of non-synonymous mutation is higher for ORF8 and ORF10 while the opposite is true for nsp1 and ORF6. We found almost similar rates of synonymous and non-synonymous case of nsp2 nsp4 and geneS. Thus, the rate of synonymous or non-synonymous mutations varies among the gene regions and for few genes the amino acid changing propensity is relatively higher than synonymous changes. For all these genes the dN/dS values are not significantly higher than 1 and the notion of purifying selection was found.

Nucleotide diversity, dS and dN values were also calculated for group B and sub-groups C, D and E (Figure1 of supplementary information) and we have identified several protein coding regions of consistent changes. These are nsp2, nsp3, nsp4, nsp6, nsp12, nsp13, nsp14, geneS, ORF3a, ORF8 and geneN. Thus, these regions are consistently evolving in SARS-CoV-2 in different lineages and may generate multiple variants. The two recently emerged groups B and E showed higher divergence of nsp9 and ORF7a respectively and mostly by amino acid changes.

### Structural mutations are associated with the viral groups

We next wanted to identify the structural mutations in all 26 protein-coding regions and map them individually onto the genome level phylogenetic tree (Figure2 of supplementary information). For this we utilized the TimeZone software on individual genes for alignment, phylogenetic analysis and tracking of structural mutations. As mentioned earlier, the protein-coding regions nsp1, nsp7, nsp8, nsp11, geneE, ORF6 and ORF7b do not experience any structural changes and are maintaining their ancient structure. Among them the small, integral membrane protein E is important at different stages of the life cycle of this virus, viz. assembly, budding, envelope formation, and pathogenesis. The highly conserved nature of this protein indicates that a specific structural requirement is necessary for the proper functioning of this protein in the viral life cycle as well as in its pathogenicity.

Several structural mutations were detected for nsp3 (proteinase) (Figure2 of supplementary information) and structural glycoprotein (S) (Figure 4). Apart from these two proteins nsp2, nsp4, RdRp, nsp13 (helicase), ORF3a and geneN also showed a considerable number of structural changes (Figure 2 of supplementary information). The mapping of all these structural mutations on to the genome-based phylogenetic tree revealed the association of several of these mutations with different groups as shown in Figure 1 and 2.

**Figure 4.**
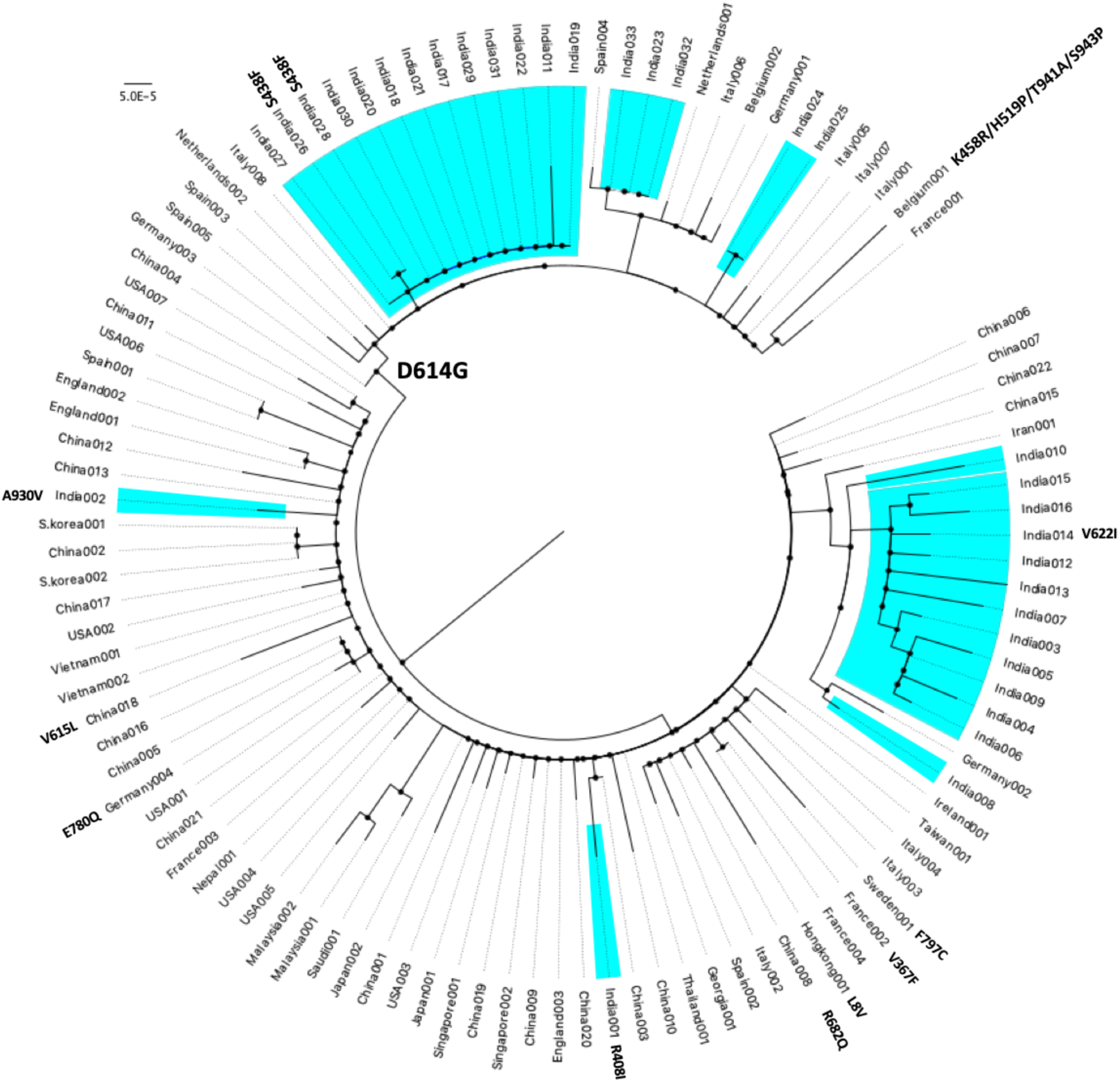
Mapping of structural mutations for spike glycoprotein (S) onto the genome-based phylogeny.

The association of mutation is based on the four genes: nsp2, nsp12 (RdRp), geneS and ORF8 (Table 1). The amino acid change V198I in nsp2 is associated with the emergence of group E, comprising mostly of Indian isolates and those from Germany and Iran. The change P323L in the RdRp gene is a signature for all the members of group B except isolate Germany003. This group is also associated with the change of D614G in spike glycoprotein S. This particular group has probably emerged in the later phase of the pandemic and harbour a considerable amount of Indian isolates. For group D, the change L84S in ORF8 is present in all of its members. Thus in comparison to the initial isolates of group C, all other groups has a unique signature of mutations, which can describe the phylogenic position of any isolate.

**Table 1.** Various phylogenetic groups and associated structural changes.

| Group | Protein coding region | Amino acid changes |
| --- | --- | --- |
| B | nsp12 (RdRp)* | P323L |
| B | Structural glycoprotein (S) | D614G |
| D | ORF8 | L84S |
| E | nsp2 | V198I |

\* except isolate Germany003

Our study revealed three protein coding regions: nsp8, nsp11 and ORF7b to be conserved throughout all the genomes we have considered. Among these gene products, nsp8 forms a complex with nsp7 which can work as multimeric RNA polymerase in order to extend primer for RdRp action (te Velthuis et al., 2012). The diversity analysis in this study clearly indicates the conservation of such complex with no structural changes in both nsp8 and nsp7 (Figure 3). Thus both of these partner proteins are highly conserved probably due to the proper complex formation for RdRp activity. On the other hand, there are seven proteins experiencing structural changes in all four phylogenetic groups, these proteins are nsp3, nsp12 (RdRp), nsp13 (helicase), structural glycoprotein (S), ORF3a, ORF8 and nucleocapsid phosphoprotein (N). Among these, the structural mutations in nsp12 and spike glycoprotein S are group specific and we have further studied that how these structural changes can potentially modulate its function.

All these proteins are important for viral life-cycle and pathogenesis and the structural changes among them may manifest phenotypic alterations and in turn change the viral pathogenesis. With rigorous computational analysis we have further tried to evaluate the impact of these structural changes in spike glycoprotein functionality. It is important to note that many of the mutations identified by our study are present in single viral isolate, possibly this can also happen either due to presence of limited number of viral isolates in our study or due to sequencing errors. Whereas, the cardinal mutations defining different groups are present in multiple viral isolates from different countries, collected in different time points.

### Spike protein functionality predicted to be affected by structural mutations

It is well established that spike (S) protein plays a pivotal role during viral entry into the host cell (Gordon et al., 2020; Ou et al., 2020). The viral entry into the host cells is mediated by the initial binding of the receptor-binding domain (RBD) present at S1 subunit of the S protein to the host ACE2 receptor followed by fusion of the viral spike and host membrane through S2 subunit located on the surface of the S protein. RBD consists of 331-524 amino acid residues of the SARS-CoV-2 S protein (Tai et al., 2020). In comparison with the S protein sequence pattern from index genome, Indian spike protein sequences possess five different variants arising due to 5 different structural mutations. Two of the protein variants are due to mutations present in the RBD at 408th (R408I) and 438th (S438F) position. Thus, these two mutations may directly affect the structural conformation required for the receptor binding. The other three mutations (D614G, V622I and A930V) found on the Indian specific SARS-CoV-2 S proteins are present outside the RBD.

The mutation R408I at the RBD involved positively charged arginine (R) with hydrophilic basic side chain is mutated to non-polar hydrophobic isoleucine (Ile) amino acid with C-beta branch. Structurally, the guanidine moiety with three amino groups has been replaced with the small methyl substitution in this mutation (R408I). As a result, the non-polar hydrophobic side chains present in Ile act as hindrance towards the accessible aqueous environment. Also, arginine with its long side-chain has greater flexibility in comparison to the mutated Ile with less to moderate side-chain flexibility which can affect the host cell bindings. The basic characteristics of these two proteins are so unique that their exchange through mutation will result in conformational change in the structure which may contribute towards the host cell recognition for receptor binding. According to our secondary structure prediction of India specific sequences with respect to the index S sequence, we observed a change in secondary structural conformation. In the 408 region of the index S protein possess a helices conformation which is changed into sheet like structures due to mutation in the Indian isolated S protein. Apart from this, tertiary structure analysis also shows changes in the structural conformation in RBD due to this mutation. The protein conformational space has been reduced from 131 ml/mol to the 121 ml/mol as the bulky terminal guanidine moiety in arginine is replaced by a small alkyl group. As a consequence, the electrostatic forces also get modified. The final protein binding conformational enthalpy (ΔΔG) is 0.133 kcal/mol after mutation suggests much stabilized conformations. Change in vibrational entropy (ΔΔS_Vib_ ENCoM: 0.178 kcal.mol^-1^.K^-1^) suggest increase in the molecular flexibility of the protein backbones which might affect the interactions with the host cell receptors. A change in inter-atomic interaction is calculated for both wild type and mutated type. In both the conformations both Arg408 and Ile408 were able to form stable intra-hydrogen bond interactions with Glu406 and Ile410, whereas ionic interaction between Arg408 and Gln414 is absent after the mutations (Figure 5A, 5B).

**Figure 5.**
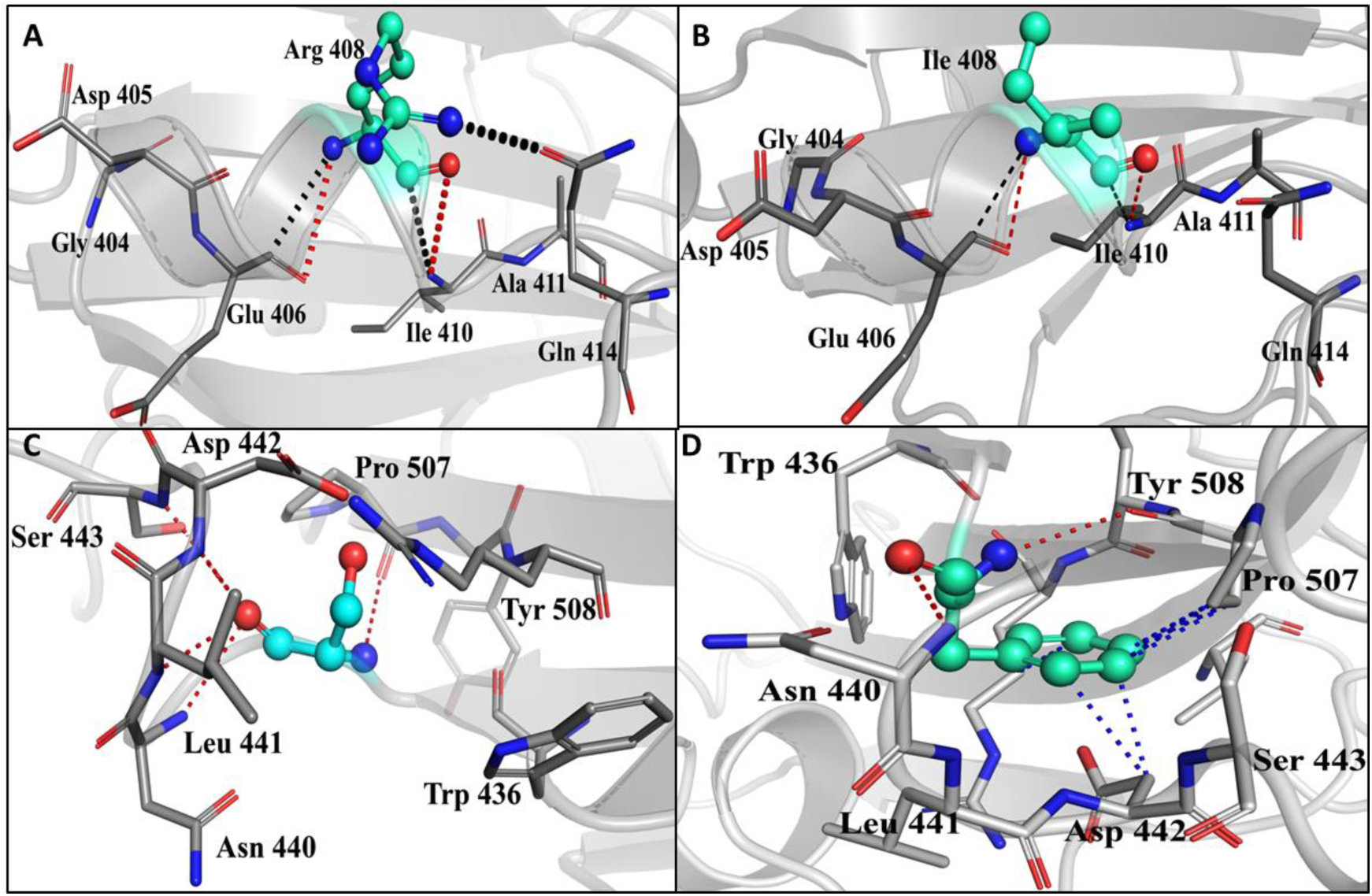
Comparison of the inter-atomic interaction in the wild type and mutated form. (A, B): In both the conformations the amino acid positions Arg408 and Ile408 were able to form stable intra-hydrogen bond interactions between Glu406 and Ile410, whereas ionic interaction between Arg408 and Gln414 is absent after the mutations. (C, D): The mutation S438F, resulted in the loss of ionic interactions (Asn440) and the hydrogen bonding formation. All the neighboring residues are depicted in gray stick representations. Significant residues in wild type and mutated type are indicated by cyan color ball and stick representations. All the intra hydrogen bonds with the neighboring residues are shown in red dotted lines whereas black colour dotted lines signify vdw interactions and hydrophobic interactions are represented by blue dotted lines.

Another mutation S438F at the RBD of the S1 subunit was observed in the Indian isolates India026 and India028 specific S protein sequence in comparison to that of the index sequences. The polar uncharged residue, serine (S) is mutated to non-polar aromatic phenylalanine (F). Similar to the previous mutations, due to this exchange it loses the hydrophilic polar nature of the hydroxyl (-OH) moiety and attains a hydrophobic aromatic side chain that can repel themselves away from the aqueous environment. Interestingly, this replacement leads to gain in the flexibility which has an impact on their secondary structure. Our analyses suggest that the β-turn like coil conformations on the Wuhan isolated structures is converted onto the more flexible sheet-like conformations at 438 position. In tertiary structure analysis, it is observed that since a hydroxyl group has been substituted by phenyl substituent, more protein conformational space is occupied. As a result, the molar volume is increased from 67.8 ml/mol to the 121.1 ml/mol. Expectedly, the electrostatic and binding entropies (ΔΔG) have been modified to the 1.942 kcal/mol which signify more stabilized conformations. The introduction of bulky phenyl groups led to the change in vibrational entropy (ΔΔS_Vib_ ENCoM: −0.545 kcal.mol^-1^.K^-1^), which implies lower molecular flexibility of the backbone protein. The mutation also has impact on the structural integrity. In the wild type structure, the intramolecular hydrogen bonding has been observed between Ser438 to the Leu441, Asp442, Ser443 and Pro507. On mutation, there is a loss of ionic interactions and the hydrogen bonding has been modified between Phe438 to the Asn440 and Pro507 (Figure 5C, 5D). All these data suggest that the mutation S438F might affect the bindings of the S protein with the host receptor.

The S protein for a considerable number of Indian isolates experiences another mutation at the 614^th^ position (D614G) which is located outside of the RBD at the S2 domain for entire group B isolates. The negatively charged aspartic acid (D) containing the acidic side chain is replaced by the non-polar glycine (G) residue. As a consequence hydrophilic nature and flexibility are lost, whereas the hydrophobic characteristic is acquired. As the properties of these two amino acids are different, the mutations may modify the structural conformations that can alter the functional modification towards the host cell recognition and fusion. Apart from the change in physical characteristics, our analysis suggests that this mutation does not alter their secondary structures. Both the wild type and mutated structure attain sheet like conformations in their secondary structure analysis. However, the mutation has a significant impact on the tertiary structure as the negatively charged acidic hydrophilic group is being replaced by neutral small hydrogen (hydrophobic) moiety. This residue is located on the water accessible surface at the S2 domain which is involved in the fusion of the viral and hosts cell membranes. Thus, it is logical that due to loss in acidic nature and flexibility, this mutation might implicate the S protein-mediated viral membrane fusion towards the host cell membrane. However, the interatomic hydrogen bonding with the neighboring residues remains the same after mutation (Figure 6A, 6B).

**Figure 6.**
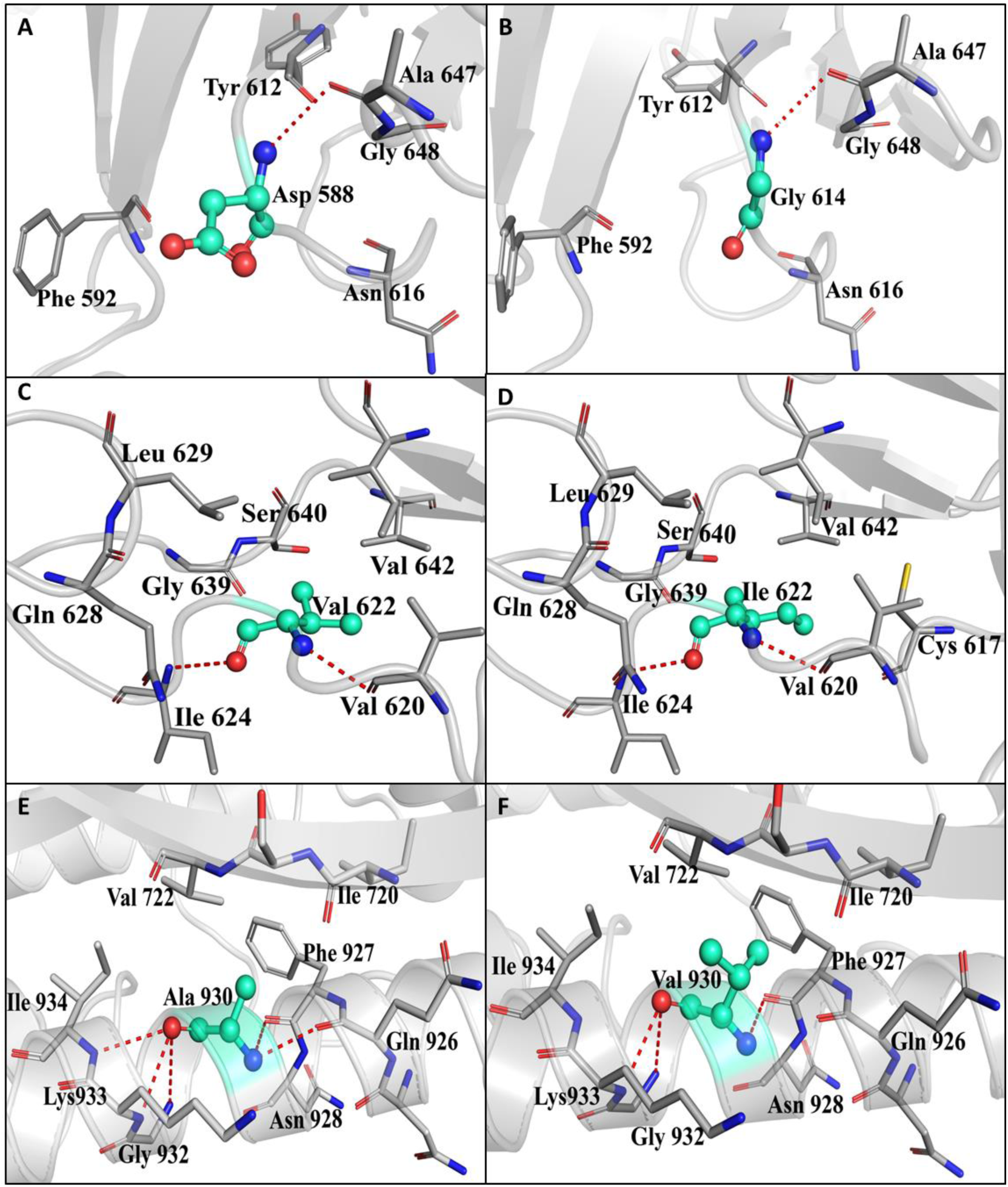
Comparison of the inter-atomic interaction in the wild type and mutated form. (A,B,C,D): The interatomic interactions with the neighboring residues remain the same after D614G and V622I mutation. (E, F): The A930V mutation affects the intra-hydrogen bond and hydrophobic interaction with the neighboring amino acids. All the neighboring residues are depicted in gray stick representations. Significant residues in wild type and mutated type are indicated by cyan color ball and stick representations. All the intra hydrogen bonds with the neighboring residues are shown in red dotted lines whereas black color dotted lines signify vdw interactions and hydrophobic interactions are represented by blue dotted lines.

Mutation at the 622^nd^ (V622I) residue is present outside of the RBD domain, specifically at the S2 subunit of the Indian isolated S protein sequence. The hydrophobic valine (V) is mutated to a similar class of residue isoleucine (I) with a greater aliphatic branch chain. As a result of this mutation changes in the vibrational entropy (ΔΔS_Vib_ ENCoM = −0.191 kcal.mol^-1^.K^-1^) were observed, which indicates a decrease in molecular flexibility causing distortion of the protein main backbones. Our analysis suggests that the mutation does not impact changes in their tertiary structure conformation. Since both valine and isoleucine have a C-beta branching headed towards the main chains, it is difficult to attain α-helical conformations by twisting as suggested by the change in electrostatic binding energies (ΔΔG = 0.786 kcal/mol). The interatomic interactions with the neighboring residues remain the same after mutation (Figure 6C, 6D). This restricted conformation can deform the neighboring local backbone of the protein which might affect the SARS-CoV-2 spike protein-mediated viral fusion with the host cell membrane.

We observed another mutation (A930V) in the Indian isolated spike sequence at the S2 subunit. The hydrophobic aliphatic alanine (A) is mutated to valine (V) with similar hydrophobic characteristics. However, the valine contains a bulkier C-beta branched side chain that is bulkier with lesser flexibility. As a result of bulkier C-branching in valine the main skeletal backbone protein resists itself for twisting to attain restricted α-helical conformations, which is validated through the analysis of secondary structure prediction (Figure3 of supplementary information). Whereas, the helix like conformation of alanine turns into a stable flat extended sheet like structure. This stable conformational change is confirmed by binding conformational enthalpy change (ΔΔG = 0.636 kcal/mol). The vibrational entropy changes (ΔΔSVib ENCoM = −0.432 kcal.mol-1.K-1) for this mutation signify decrease in molecular flexibility. The mutation affects the intra-hydrogen bond and hydrophobic interaction (Val722) with the neighboring amino acids. In the wild type protein, Ala930 makes five hydrogen bonding with Gln926, Phe937, Gly932, Lys 933 and Ile934, whereas Val930 in the mutated protein makes three hydrogen bonds with Phe927, Gly932 and Lys933 (Figure 6E, 6F). As this mutation is present on the surface of the S2 subunit of the S protein, it may influence the spike protein driven fusion of the SARS-CoV-2 and host membrane.

### RdRp protein functionality also predicted to be affected by structural mutations

The SARS-CoV-2 nsp12 is RNA dependent RNA polymerase (RdRp) consists of 932 amino acids which are located in the polyprotein, from 4393 – 5324 aa. Structurally, SARS-CoV-2 nsp12 protein is categorized into N-terminal (1-397 aa) and a polymerase domain (398-919 aa) and is comparable to the previous SARS-CoV nsp12 (PDB ID: 6NUR). The polymerase domain can be subdivided into structurally different three subunits; a finger (398–581 and 628–687 aa), a palm (582–627 and 688–815 aa), and a thumb subunit (816–919 aa) which adopts a “right-handed” cupped shaped conformation (Yin et al., 2020). The active site of RdRp is present at the interface between finger and thumb subunits of the SARS-CoV-2 RdRp, which is a conserved domain with MERS and SARS-nsp12. The active site is situated in the center of the substrate domain where RNA synthesis takes place on the entry of the RNA template from template entry channel and NTP (Nucleoside Triphosphate) through the NTP entry channel (Peersen OB., 2017). The structural analysis of the SARS-CoV-2 specified nsp12 revealed the presence of seven conserved motifs (A-G motifs) (Figure 7). These motifs play crucial roles towards the binding of template and nucleotide as well as catalysis of the NTP addition (Poch et al., 1989; Bruenn JA., 2003; Mirza et al., 2019). Several antiviral drugs act by attaching onto the first replicated base pairs and terminates the chain elongation process. The RdRp binding affinity towards the template RNA has been increased by the association of the co-factor nsp7 and nsp8 with the nsp12. RNA polymerization activity is found to be inhibited in presence of the active triphosphate form of Remdesivir (RTP), which is under clinical trial (Yin et al., 2020; Shannon et al., 2020).

**Figure 7:**
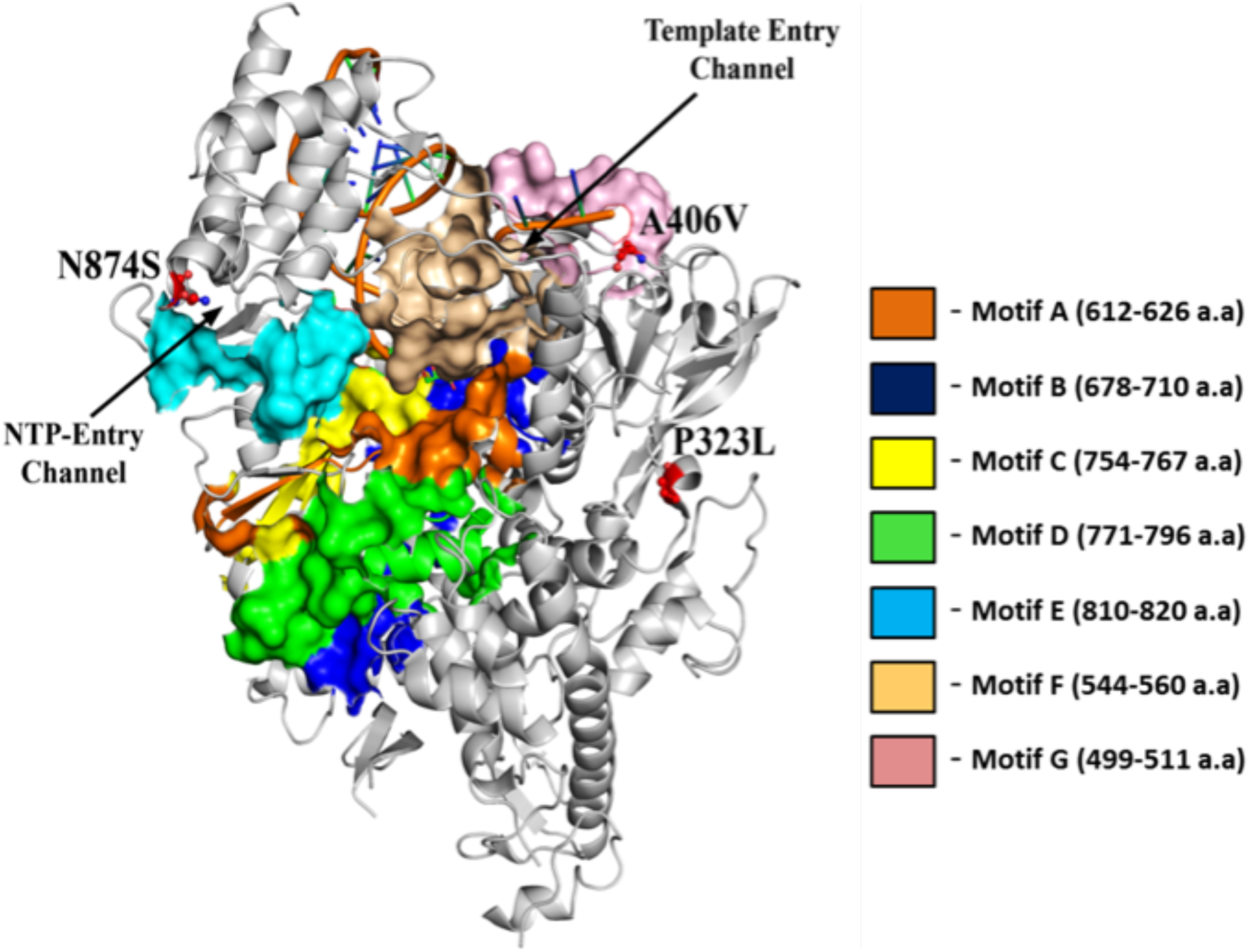
Cartoon representation of SARS-CoV-2 RdRp (PDB ID: 7BV2) with the spatial arrangement of various structural motif A-G. RdRp protein is complexed with an RNA template. RdRp structure is depicted with the white cartoon representations with several structural motifs. The motifs are colored as follows A-Orange; B-Blue; C-Yellow; D-Green; E-Cyan; F-light yellow; G-pink. The template RNA is portrayed in cartoon representation with rainbow color.

The mutation P323L in the RdRp gene which is a signature of all the members of group B is present in the N-terminal (1-397 aa) region of the SARS-CoV-2 RdRp protein. Previous studies revealed that proper folding of the neighboring RdRp’s finger subdomain (397-581 aa & 628-687 aa) and proper positioning of the polymerase active site is fully depended on the N-terminal structural conformations. In this mutation a hydrophobic proline (P) has been mutated to the similar hydrophobic aliphatic leucine (L). Secondary structure analysis revealed the structural conformation changes after mutation. Although proline is known as “helix breakers” since they interrupt the stability of the α-helical conformations of the backbone protein but in this case proline is present in β-turn-like conformations (Figure4 of supplementary information). Whereas mutated residue leucine contains the bulkier C-beta branched aliphatic side chains and found present as a sheet-like conformation (Figure4 of supplementary information). In the tertiary structure analysis suggest that the mutation is present in the N-terminal region which is connected to the finger subdomain (Figure 7). So this mutation on the N-terminal domain may hamper the proper folding of the SARS-CoV-2 RdRp backbone protein, which can play a vital role for the protein’s consequent degradation (te Velthuis AJ., 2010).

In comparison to the Wuhan strain, Indian isolated SARS-CoV-2 specified RdRp experienced two other mutations (A406V and N874S) as shown in Figure 7. Alanine 406 is not in the active site domain but this is present on the template RNA entry channel. In this mutation hydrophobic aliphatic alanine is converted to a similar class of non-polar valine residue. As a result such mutation is not expected to cause any significant change in protein functionalities. However, due to the presence of the bulkier C-beta branched-chain, it restricted the protein main backbone to attain the alpha-helical conformations. Also, the heavy C-beta branching reduces the side chain flexibilities, which can modify the RNA template entry and if the RNA template cant enters on to the catalytic domain, RNA synthesis finally gets hampered. In another mutation (N874S) the polar uncharged asparagine (N) is replaced with similar polar uncharged serine (S). However, there is a different polar group with Asn carry an amide group and Ser has a hydroxyl group. Also, due to a slight reduction of side-chain length in Ser as compared to Asn, flexibilities of the β-turn conformation will be hampered. As this mutation is observed near the NTP entry channel (Figure 7), any changes in the conserved β-turn conformation of the backbone protein might affect the NTP channel and interfere the RNA synthesis. Interestingly, such Asn to Ser mutation has been found to confer oseltamivir resistance to H5N1 influenza A viruses (Kiso et al., 2011).

## Conclusion

In this study, we analyzed the whole genome as well as all the protein coding regions of 109 genomes of SARS-CoV-2 considered from different countries of higher incidence of COVID-19. SARS-CoV-2 is the pathogen that causes severe pneumonia in several of the affected individuals and is responsible for about 0.2 million deaths worldwide. With the advent of high-throughput next generation sequencing technology the researchers around the world have access to huge number of SARS-CoV-2 genome sequences. We also took the opportunity of the availability of this resource in order to study the genome divergence pattern of this pathogenic virus for identification of emerging strains with distinct genotypes. This study has potential for better understanding of the evolution and transmission of SARS-CoV-2 genome globally, locally and even within a single host. Although our dataset comprises of 109 genomes, we considered the isolates from various countries of the world with evidence of early viral transmission initially originating from China and with emphasis on India which is in its early stages of COVID-19 pandemic.

Our results describe the clear association of four structural changes with that of group structure from phylogenomics analysis of the SARS-CoV-2 isolates in this present pandemic situation. Also the structural studies performed here indicate the probable connection of these variants with varied level of virulence potentials of SARS-CoV-2. In case of India, this study with considerable number of genome sequences portrays to a certain extent the population structure of COVID-19. Further, it is apparent that multiple emergent variants of viral genomes are circulating in Indian and new genome sequences can be characterised in this way. Obviously more sequencing of viral genomes and associated clinical outcome data may further validate the relevance of our findings and implicate the viral phenotype with group structure. This prospect has tremendous impact on public health in order to mitigate the present pandemic situation.

## Author Contributions

AT and SP2 conceived and designed the experiments. DB, SP1, AL and SP2 performed the experiments. DB, SP1, AT and SP2 analysed the data. DB, SP1, AT and SP2 wrote the manuscript.

## Conflict of Interest Statement

The authors declare that the research was conducted in the absence of any commercial or financial relationships that could be construed as a potential conflict of interest.

## Acknowledgement

DB and SP1 have received Senior Research Fellowship (SRF) from Indian Council of Medical Research. AL has received Senior Research Fellowship (SRF) from Council of Scientific and Industrial Research. SP2 has received the Ramanujan Fellowship from SERB, Govt. of India.

## Notes

### Competing Interest Statement

The authors have declared no competing interest.

