## Supplementary Figure 1 for "Emergence of multiple variants of SARS-CoV-2 with signature structural changes"

Supplementary Figure 1: Average nucleotide diversity (pi), rate of synonymous (dS) and non-synonymous (dN) substitution rates for four phylogenetic groups (B, C, D & E)

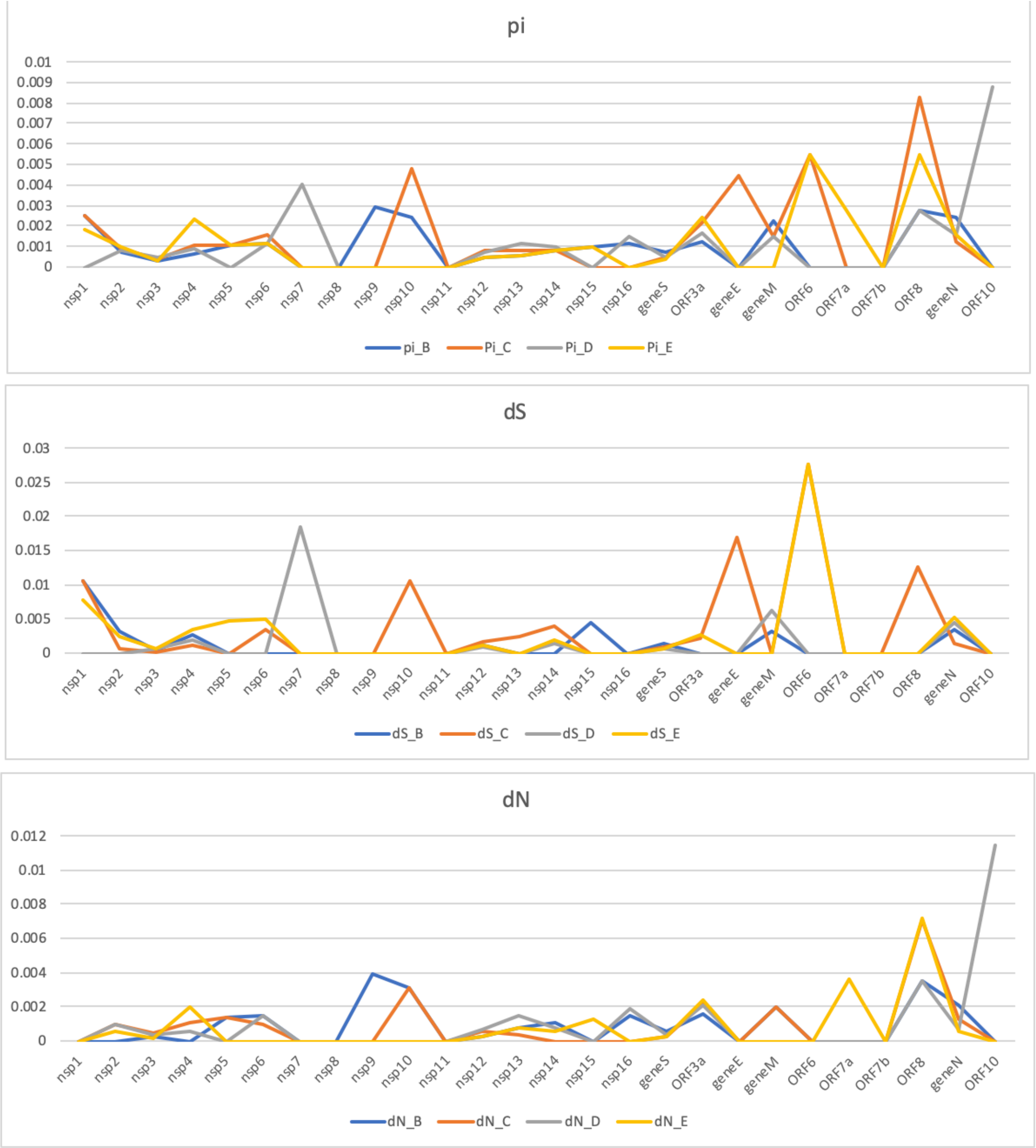
