## Supplementary Figure 2 for "Emergence of multiple variants of SARS-CoV-2 with signature structural changes"

Supplementary Figure3. Mapping of structural mutations in whole genome phylogenetic tree. For isolates from India blue background was used.

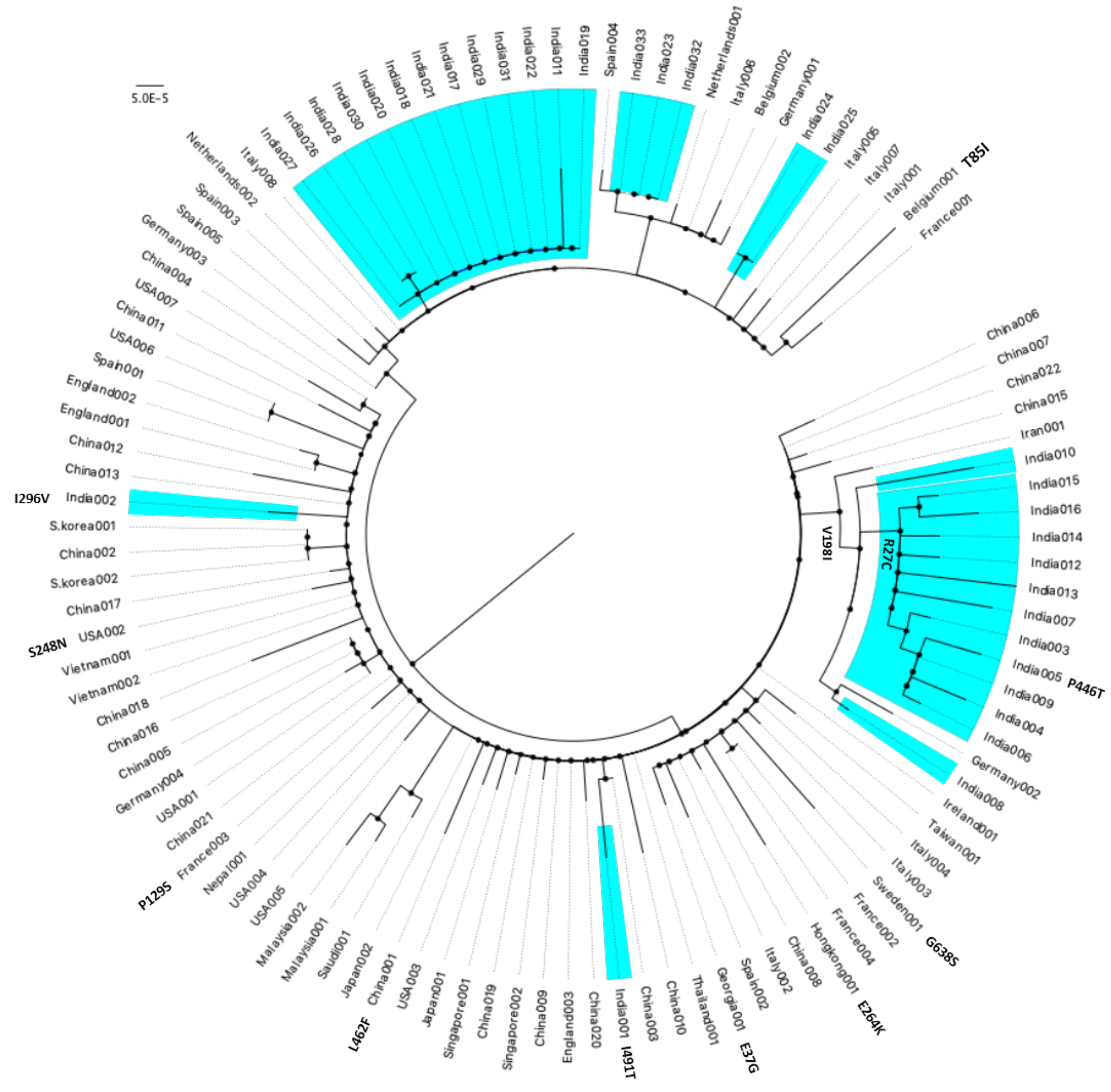


**nsp2**

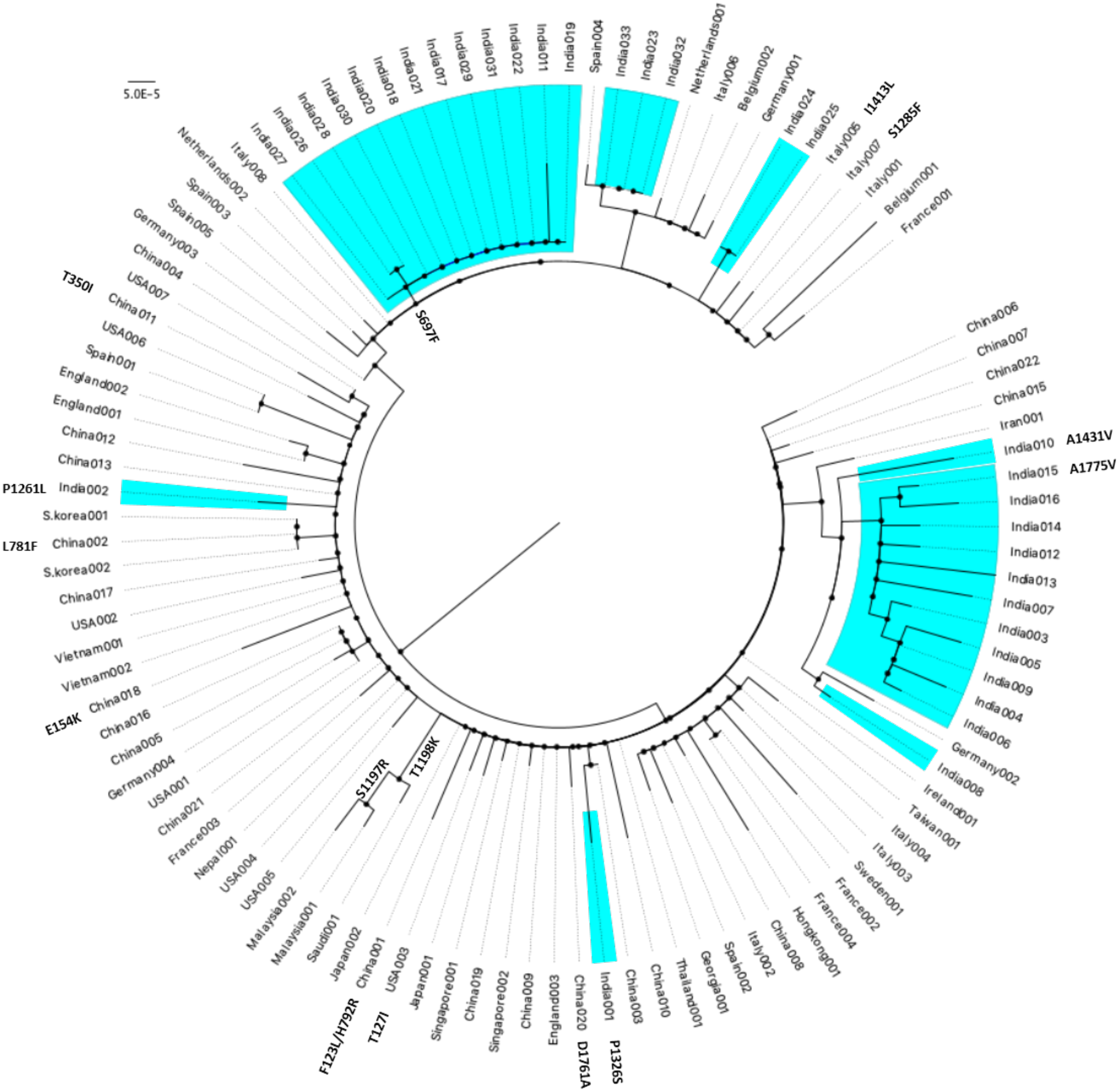

5.0E-5
India019
India011
India022
India031
India029
India017
India021
India018
India020
India030
India028
India026
India027
Italy008
Netherlands002
Spain003
Spain005
Germany003
China004
USA007
T350I
China011
USA006
Spain001
England002
England001
China012
China013
P1261L
India002
S.korea001
L781F
China002
S.korea002
China017
USA002
Vietnam001
Vietnam002
E154K
China018
China016
China005
Germany004
USA001
China021
France003
Nepal001
USA004
USA005
Malaysia002
Malaysia001
Saudi001
Japan002
China001
USA003
F123L/H792R
T127I
Japan001
Singapore001
China019
Singapore002
China009
England003
China020
D1761A
India001
P1326S
China003
China010
Thailand001
Georgia001
Spain002
Italy002
China008
Hongkong001
France004
France002
Sweden001
Italy003
Italy004
Taiwan001
Ireland001
India008
Germany002
India006
India004
India009
India005
India003
India007
India013
India012
India014
India016
India015
A1775V
India010
A1431V
Iran001
China015
China022
China007
China006
France001
Belgium001
Italy001
Italy007
S1285F
Italy005
India025
I1413L
India024
Germany001
Belgium002
Italy006
Netherlands001
India032
India023
India033
Spain004
S697F
S1197R
T1198K


**nsp3**

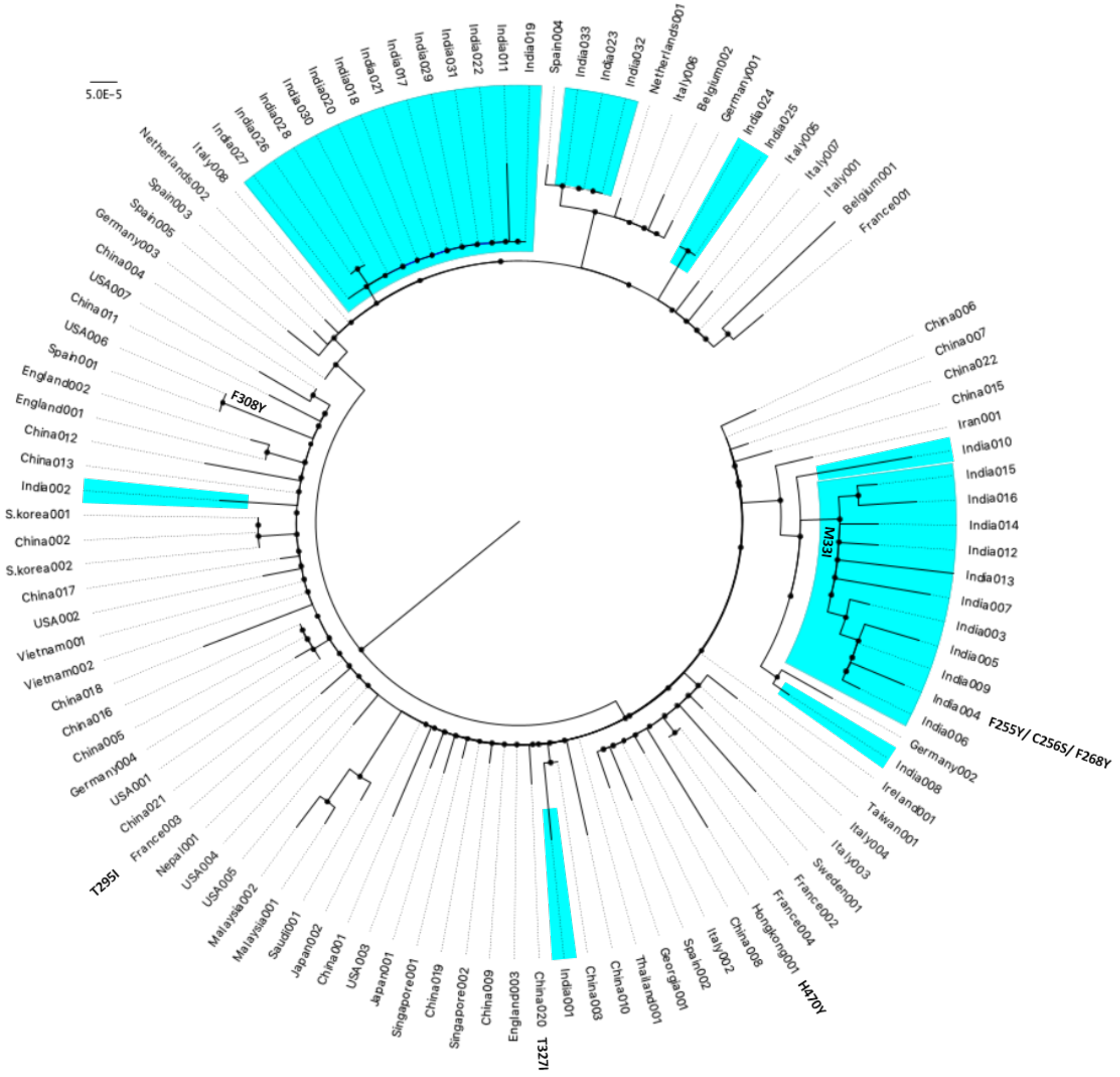
5.0E-5
Netherlands002
Italy008
India027
India026
India028
India030
India020
India018
India021
India017
India029
India031
India022
India011
India019
Spain004
India033
India023
India032
Netherlands001
Italy006
Belgium002
Germany001
India024
India025
Italy005
Italy007
Italy001
Belgium001
France001
China006
China007
China022
China015
Iran001
India010
India015
India016
India014
India012
India013
India007
India003
India005
India009
India004
India006
Germany002
India008
Ireland001
Taiwan001
Italy004
Italy003
Sweden001
France002
France004
Hongkong001
China008
Italy002
Spain002
Georgia001
Thailand001
China010
China003
India001
China020
England003
China009
Singapore002
China019
Singapore001
Japan001
USA003
China001
Japan002
Saudi001
Malaysia001
Malaysia002
USA005
USA004
Nepal001
France003
China021
USA001
Germany004
China005
China016
China018
Vietnam002
Vietnam001
USA002
China017
S.korea002
China002
S.korea001
India002
China013
China012
England001
England002
Spain001
USA006
China011
USA007
China004
Germany003
Spain005
Spain003
F308Y
M33I
F255Y/ C256S/ F268Y
H470Y
T327I
T295I

**nsp4**

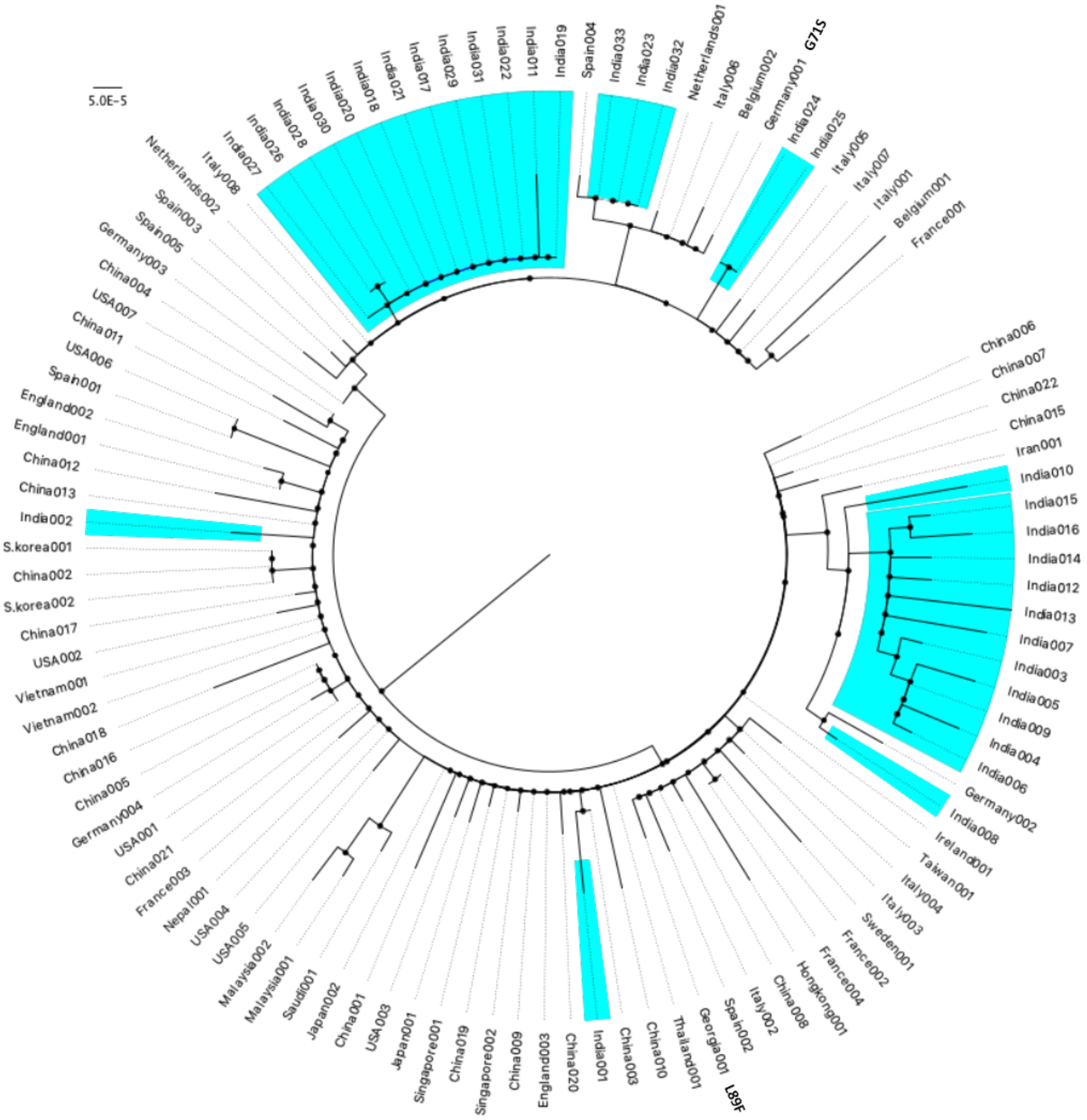
5.0E-5
Netherlands002
Italy008
India027
India026
India028
India030
India020
India018
India021
India017
India029
India031
India022
India011
India019
Spain004
India033
India023
India032
Netherlands001
Italy006
Belgium002
Germany001
G71S
India024
India025
Italy005
Italy007
Italy001
Belgium001
France001
China006
China007
China022
China015
Iran001
India010
India015
India016
India014
India012
India013
India007
India003
India005
India009
India004
India006
Germany002
India008
Ireland001
Taiwan001
Italy004
Italy003
Sweden001
France002
France004
Hongkong001
China008
Italy002
Spain002
Georgia001
L89F
Thailand001
China010
China003
India001
China020
England003
China009
Singapore002
China019
Singapore001
Japan001
USA003
China001
Japan002
Saudi001
Malaysia001
Malaysia002
USA005
USA004
Nepal001
France003
China021
USA001
Germany004
China005
China016
China018
Vietnam002
Vietnam001
USA002
China017
S.korea002
China002
S.korea001
India002
China013
China012
England001
England002
Spain001
USA006
China011
USA007
China004
Germany003
Spain005
Spain003

**nsp5**

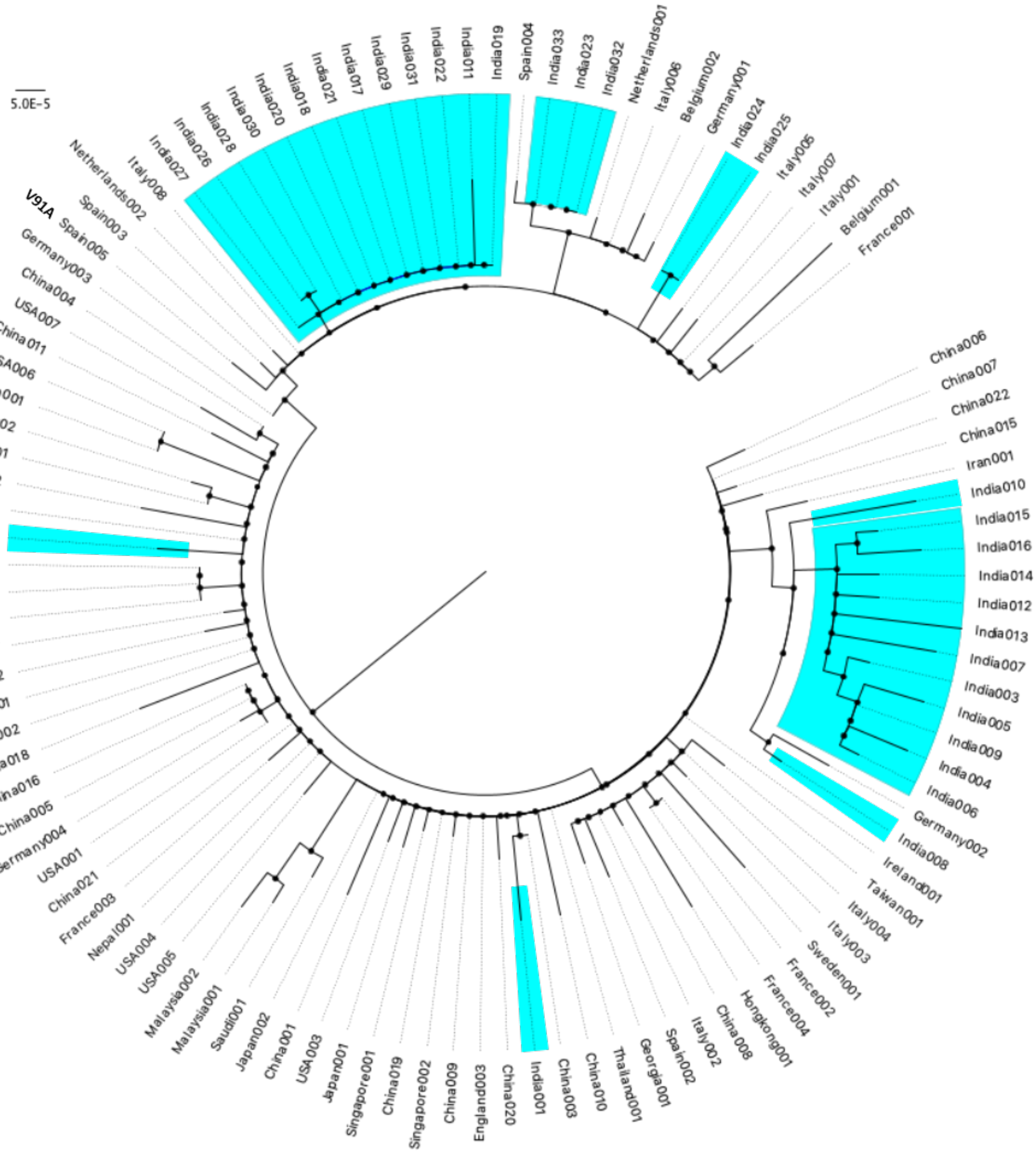
5.0E-5
V91A
India019
India011
India022
India031
India029
India017
India021
India018
India020
India030
India028
India026
India027
Italy008
Netherlands002
Spain003
Spain005
Germany003
China004
USA007
China011
USA006
Spain001
England002
England001
China012
China013
India002
S.korea001
China002
S.korea002
China017
USA002
Vietnam001
Vietnam002
China018
China016
China005
Germany004
USA001
China021
France003
Nepal001
USA004
USA005
Malaysia002
Malaysia001
Saudi001
Japan002
China001
USA003
Japan001
Singapore001
China019
Singapore002
China009
England003
China020
India001
China003
China010
Thailand001
Georgia001
Spain002
Italy002
China008
Hongkong001
France004
France002
Sweden001
Italy003
Italy004
Taiwan001
Ireland001
India008
Germany002
India006
India004
India009
India005
India003
India007
India013
India012
India014
India016
India015
India010
Iran001
China015
China022
China007
China006
France001
Belgium001
Italy001
Italy007
Italy005
India025
India024
Germany001
Belgium002
Italy006
Netherlands001
India032
India023
India033
Spain004

**nsp6**

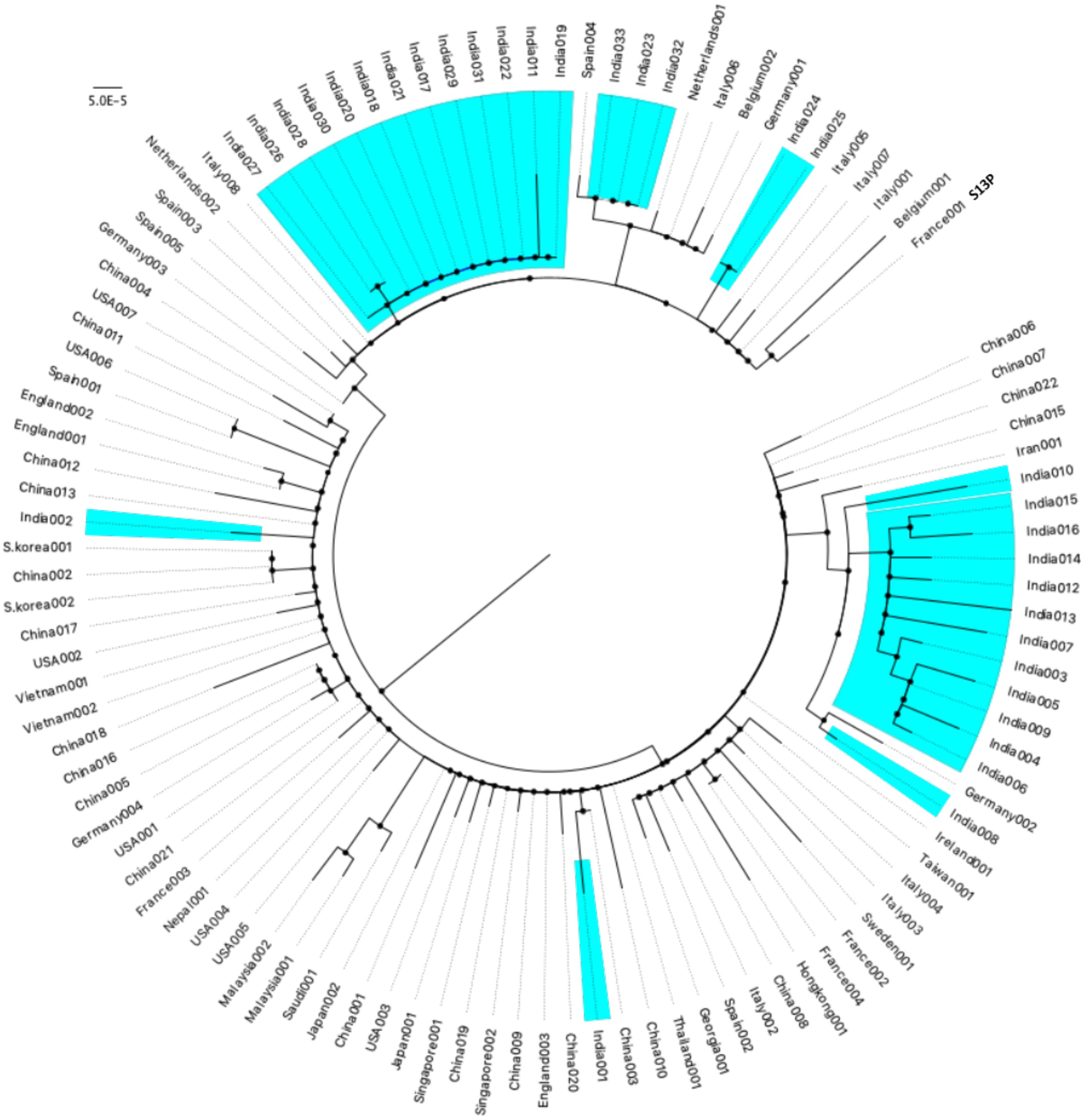
5.0E-5
Netherlands002
Italy008
India027
India026
India028
India030
India020
India018
India021
India017
India029
India031
India022
India011
India019
Spain004
India033
India023
India032
Netherlands001
Italy006
Belgium002
Germany001
India024
India025
Italy005
Italy007
Italy001
Belgium001
France001
S13P
China006
China007
China022
China015
Iran001
India010
India015
India016
India014
India012
India013
India007
India003
India005
India009
India004
India006
Germany002
India008
Ireland001
Taiwan001
Italy004
Italy003
Sweden001
France002
France004
Hongkong001
China008
Italy002
Spain002
Georgia001
Thailand001
China010
China003
India001
China020
England003
China009
Singapore002
China019
Singapore001
Japan001
USA003
China001
Japan002
Saudi001
Malaysia001
Malaysia002
USA005
USA004
Nepal001
France003
China021
USA001
Germany004
China005
China016
China018
Vietnam002
Vietnam001
USA002
China017
S.korea002
China002
S.korea001
India002
China013
China012
England001
England002
Spain001
USA006
China011
USA007
China004
Germany003
Spain005
Spain003

**nsp9**

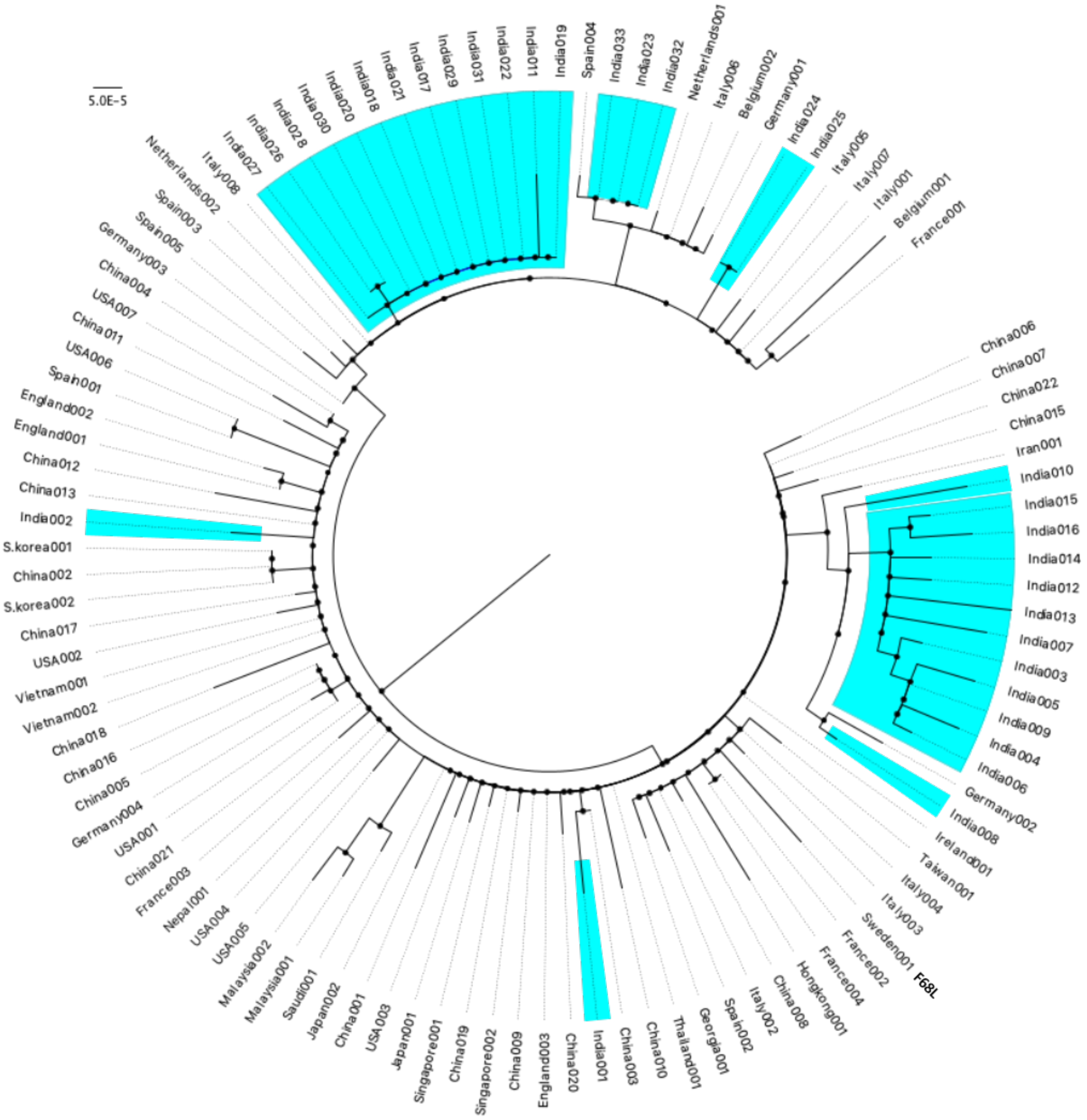
5.0E-5
India019
India011
India022
India031
India029
India017
India021
India018
India020
India030
India028
India026
India027
Italy008
Netherlands002
Spain003
Spain005
Germany003
China004
USA007
China011
USA006
Spain001
England002
England001
China012
China013
India002
S.korea001
China002
S.korea002
China017
USA002
Vietnam001
Vietnam002
China018
China016
China005
Germany004
USA001
China021
France003
Nepal001
USA004
USA005
Malaysia002
Malaysia001
Saudi001
Japan002
China001
USA003
Japan001
Singapore001
China019
Singapore002
China009
England003
China020
India001
China003
China010
Thailand001
Georgia001
Spain002
Italy002
China008
Hongkong001
France004
France002
Sweden001
F68L
Italy003
Italy004
Taiwan001
Ireland001
India008
Germany002
India006
India004
India009
India005
India003
India007
India013
India012
India014
India016
India015
India010
Iran001
China015
China022
China007
China006
France001
Belgium001
Italy001
Italy007
Italy005
India025
India024
Germany001
Belgium002
Italy006
Netherlands001
India032
India023
India033
Spain004

**nsp10**

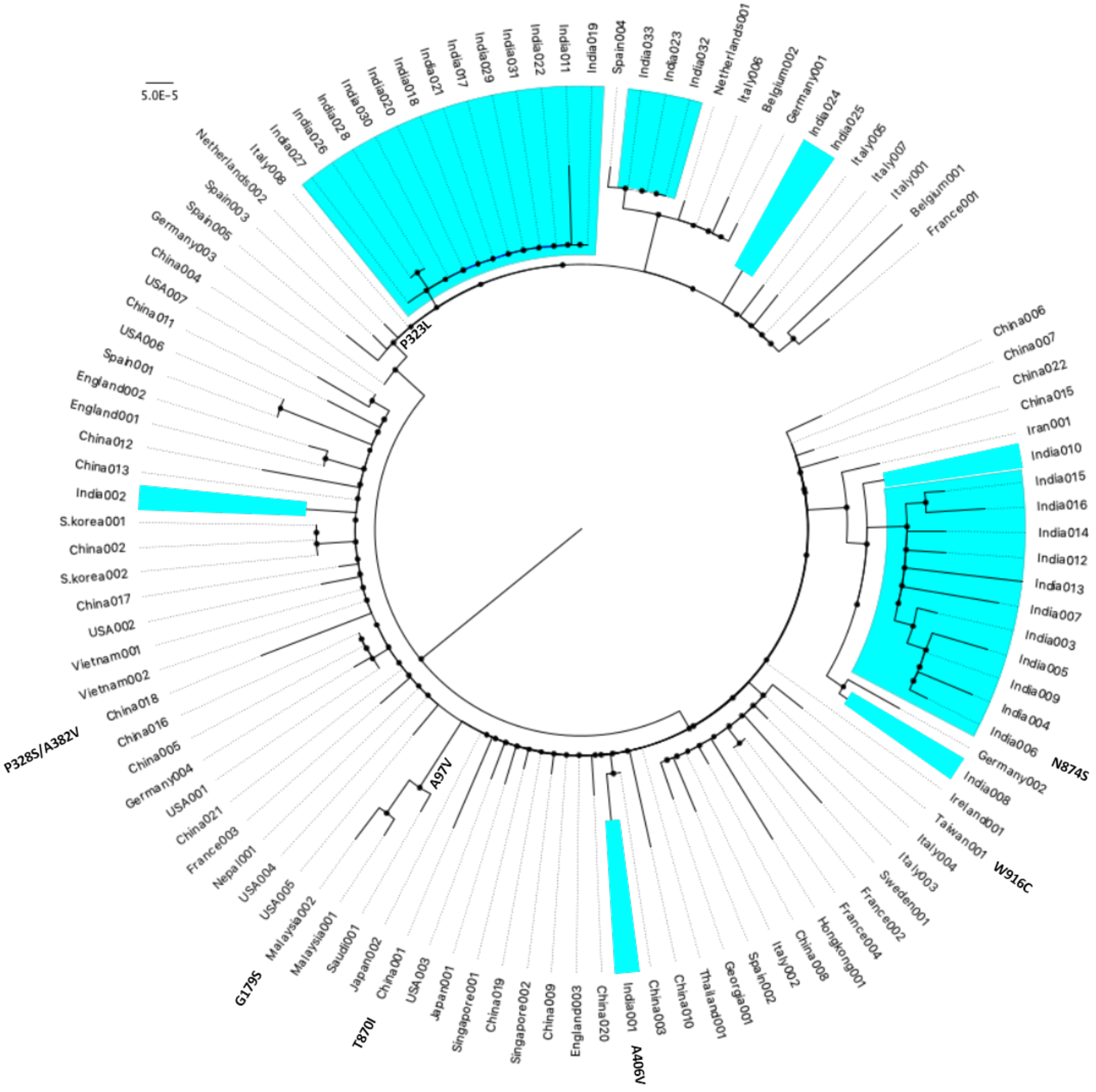
5.0E-5
Netherlands002
Italy008
India027
India026
India028
India030
India020
India018
India021
India017
India029
India031
India022
India011
India019
Spain004
India033
India023
India032
Netherlands001
Italy006
Belgium002
Germany001
India024
India025
Italy005
Italy007
Italy001
Belgium001
France001
China006
China007
China022
China015
Iran001
India010
India015
India016
India014
India012
India013
India007
India003
India005
India009
India004
India006
Germany002
India008
Ireland001
Taiwan001
Italy004
Italy003
Sweden001
France002
France004
Hongkong001
China008
Italy002
Spain002
Georgia001
Thailand001
China010
China003
India001
China020
England003
China009
Singapore002
China019
Singapore001
Japan001
USA003
China001
Japan002
Saudi001
Malaysia001
Malaysia002
USA005
USA004
Nepal001
France003
China021
USA001
Germany004
China005
China016
China018
Vietnam002
Vietnam001
USA002
China017
S.korea002
China002
S.korea001
India002
China013
China012
England001
England002
Spain001
USA006
China011
USA007
China004
Germany003
Spain005
Spain003
P323L
N874S
W916C
A406V
T870I
A97V
G179S
P328S/A382V

**nsp12**

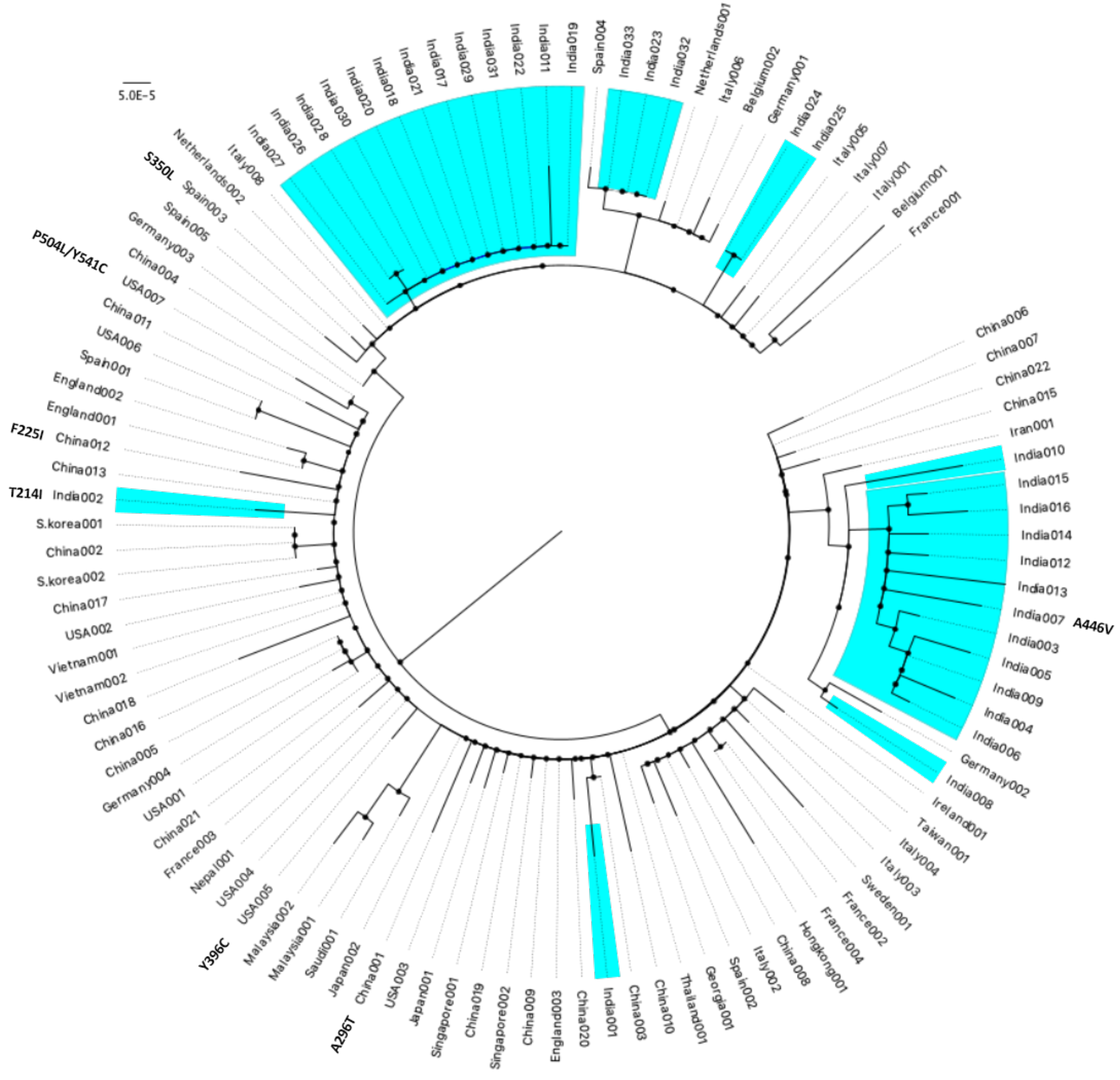

5.0E-5
Italy008
India027
India026
India028
India030
India020
India018
India021
India017
India029
India031
India022
India011
India019
Spain004
India033
India023
India032
Netherlands001
Italy006
Belgium002
Germany001
India024
India025
Italy005
Italy007
Italy001
Belgium001
France001
China006
China007
China022
China015
Iran001
India010
India015
India016
India014
India012
India013
India007
A446V
India003
India005
India009
India004
India006
Germany002
India008
Ireland001
Taiwan001
Italy004
Italy003
Sweden001
France002
France004
Hongkong001
China008
Italy002
Spain002
Georgia001
Thailand001
China010
China003
India001
China020
England003
China009
Singapore002
China019
Singapore001
Japan001
USA003
China001
A296T
Japan002
Saudi001
Malaysia001
Malaysia002
Y396C
USA005
USA004
Nepal001
France003
China021
USA001
Germany004
China005
China016
China018
Vietnam002
Vietnam001
USA002
China017
S.korea002
China002
S.korea001
T214I
India002
China013
F225I
China012
England001
England002
Spain001
USA006
China011
USA007
China004
P504L/Y541C
Germany003
Spain005
Spain003
S350L
Netherlands002


**nsp13**

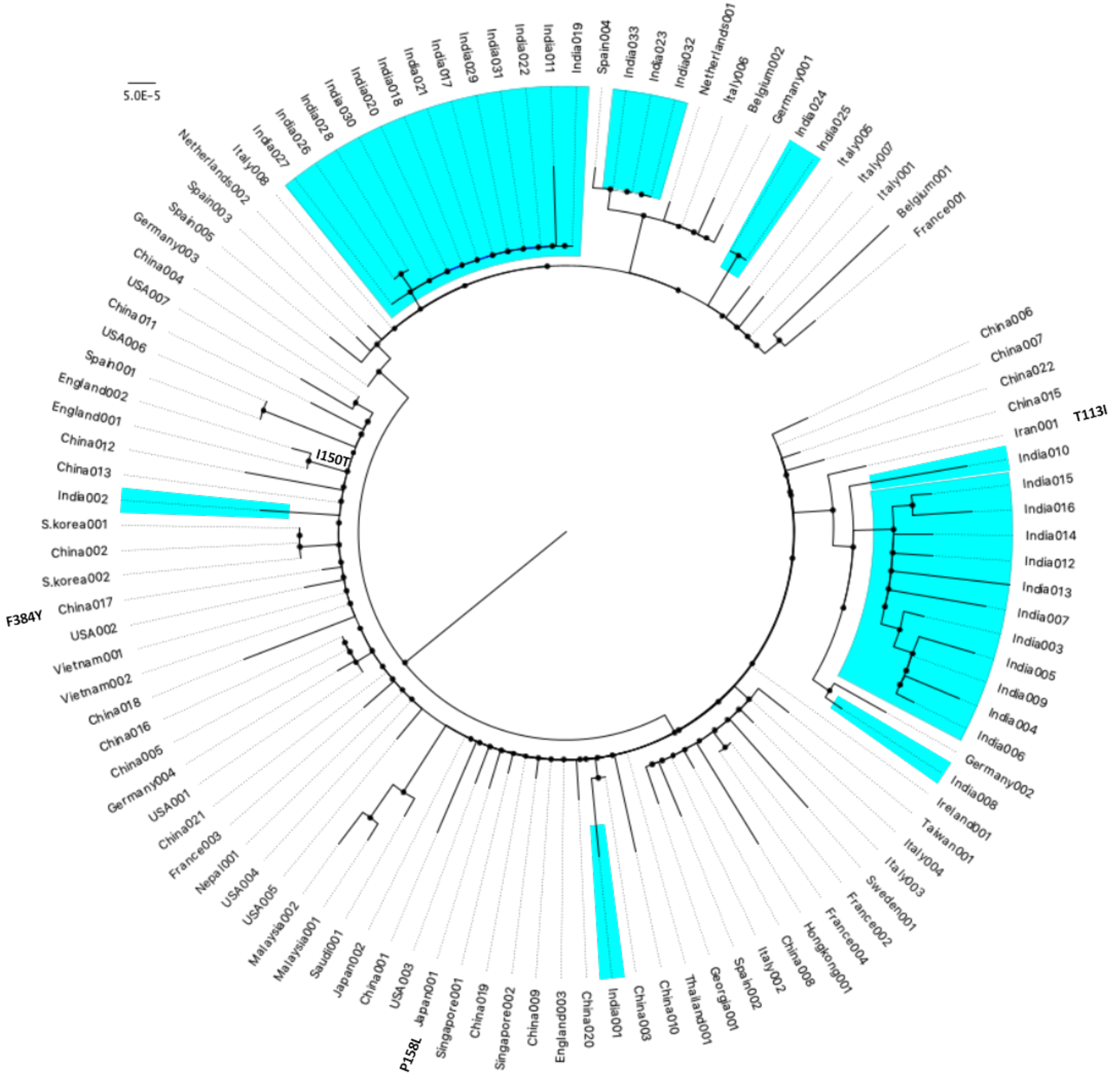
5.0E-5
Italy008
India027
India026
India028
India030
India020
India018
India021
India017
India029
India031
India022
India011
India019
Spain004
India033
India023
India032
Netherlands001
Italy006
Belgium002
Germany001
India024
India025
Italy005
Italy007
Italy001
Belgium001
France001
China006
China007
China022
China015
Iran001
T113I
India010
India015
India016
India014
India012
India013
India007
India003
India005
India009
India004
India006
Germany002
India008
Ireland001
Taiwan001
Italy004
Italy003
Sweden001
France002
France004
Hongkong001
China008
Italy002
Spain002
Georgia001
Thailand001
China010
China003
India001
China020
England003
China009
Singapore002
China019
Singapore001
Japan001
P158L
USA003
China001
Japan002
Saudi001
Malaysia001
Malaysia002
USA005
USA004
Nepal001
France003
China021
USA001
Germany004
China005
China016
China018
Vietnam002
Vietnam001
USA002
China017
F384Y
S.korea002
China002
S.korea001
India002
China013
China012
I150T
England001
England002
Spain001
USA006
China011
USA007
China004
Germany003
Spain005
Spain003
Netherlands002

**nsp14**

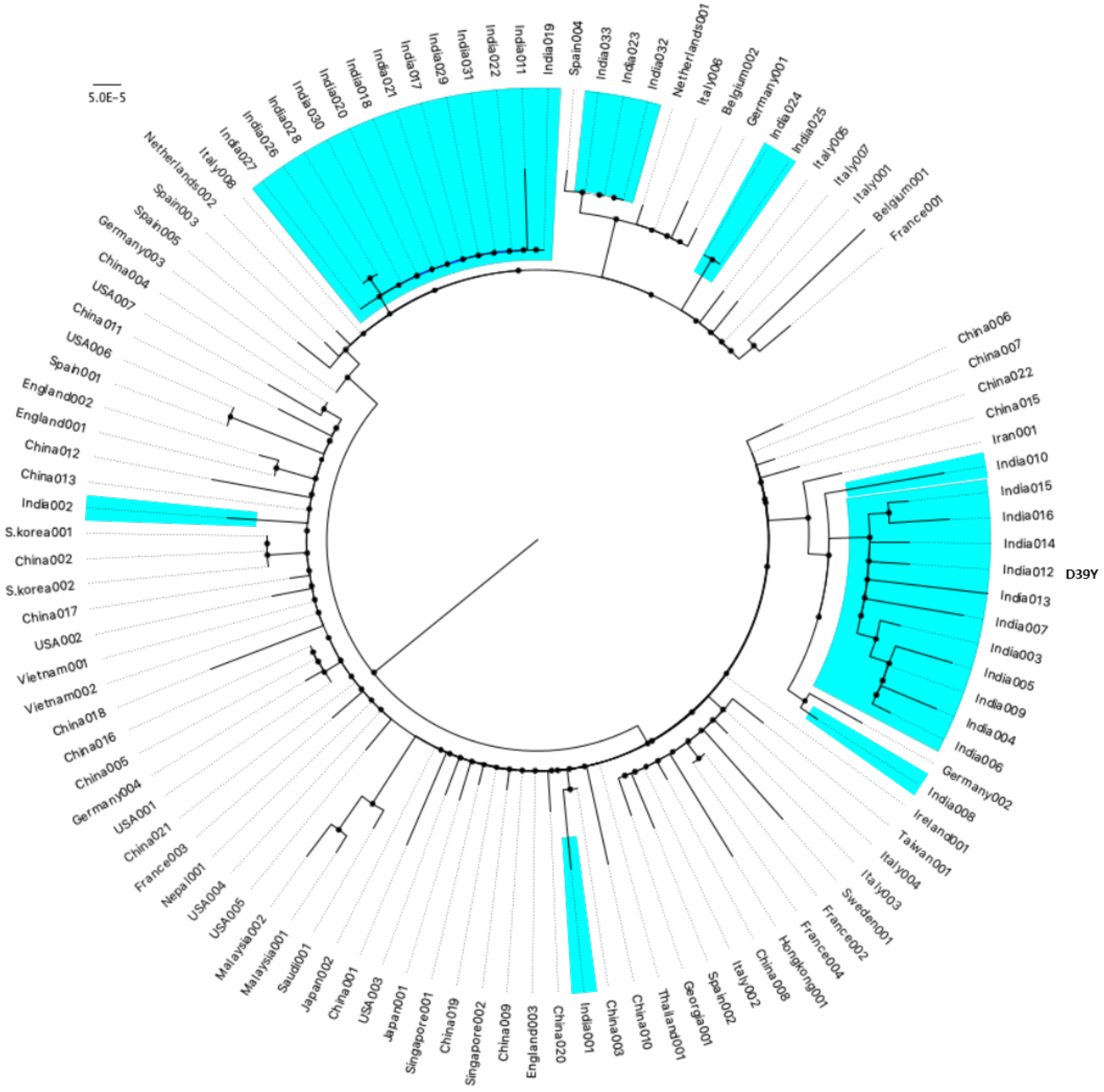


**nsp15**

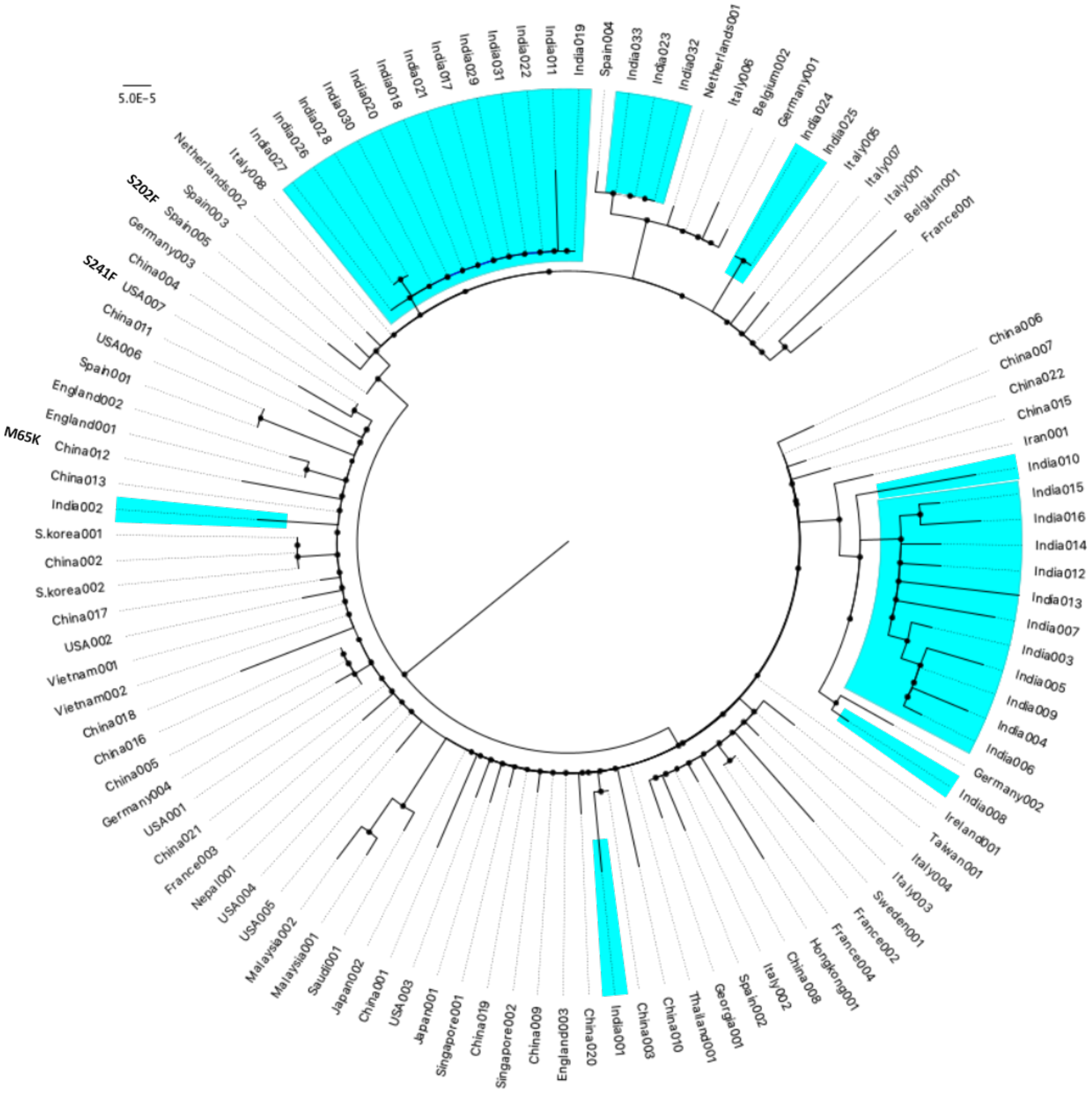

5.0E-5
S202F
S241F
M65K
Italy008
Netherlands002
Spain003
Spain005
Germany003
China004
USA007
China011
USA006
Spain001
England002
England001
China012
China013
India002
S.korea001
China002
S.korea002
China017
USA002
Vietnam001
Vietnam002
China018
China016
China005
Germany004
USA001
China021
France003
Nepal001
USA004
USA005
Malaysia002
Malaysia001
Saudi001
Japan002
China001
USA003
Japan001
Singapore001
China019
Singapore002
China009
England003
China020
India001
China003
China010
Thailand001
Georgia001
Spain002
Italy002
China008
Hongkong001
France004
France002
Sweden001
Italy003
Italy004
Taiwan001
Ireland001
India008
Germany002
India006
India004
India009
India005
India003
India007
India013
India012
India014
India016
India015
India010
Iran001
China015
China022
China007
China006
France001
Belgium001
Italy001
Italy007
Italy005
India025
India024
Germany001
Belgium002
Italy006
Netherlands001
India032
India023
India033
Spain004
India019
India011
India022
India031
India029
India017
India021
India018
India020
India030
India028
India026
India027


**nsp16**

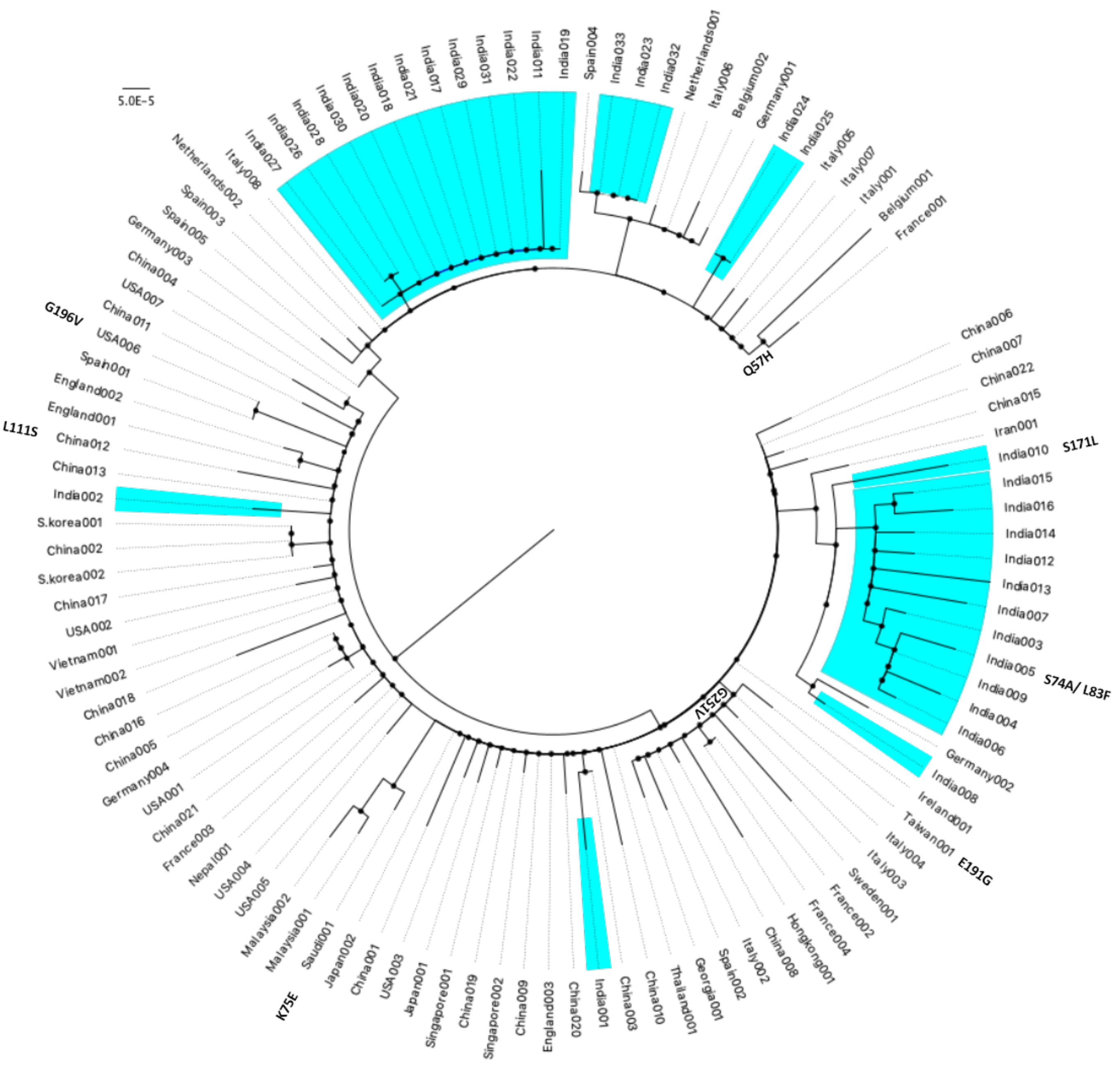
5.0E-5
India019
India011
India022
India031
India029
India017
India021
India018
India020
India030
India028
India026
India027
Italy008
Netherlands002
Spain003
Spain005
Germany003
China004
USA007
China011
USA006
Spain001
England002
England001
China012
China013
India002
S.korea001
China002
S.korea002
China017
USA002
Vietnam001
Vietnam002
China018
China016
China005
Germany004
USA001
China021
France003
Nepal001
USA004
USA005
Malaysia002
Malaysia001
Saudi001
Japan002
China001
USA003
Japan001
Singapore001
China019
Singapore002
China009
England003
China020
India001
China003
China010
Thailand001
Georgia001
Spain002
Italy002
China008
Hongkong001
France004
France002
Sweden001
Italy003
Italy004
Taiwan001
Ireland001
India008
Germany002
India006
India004
India009
India005
India003
India007
India013
India012
India014
India016
India015
India010
Iran001
China015
China022
China007
China006
France001
Belgium001
Italy001
Italy007
Italy005
India025
India024
Germany001
Belgium002
Italy006
Netherlands001
India032
India023
India033
Spain004
G196V
L111S
K75E
G251V
E191G
S74A/ L83F
S171L
Q57H

**ORF3a**

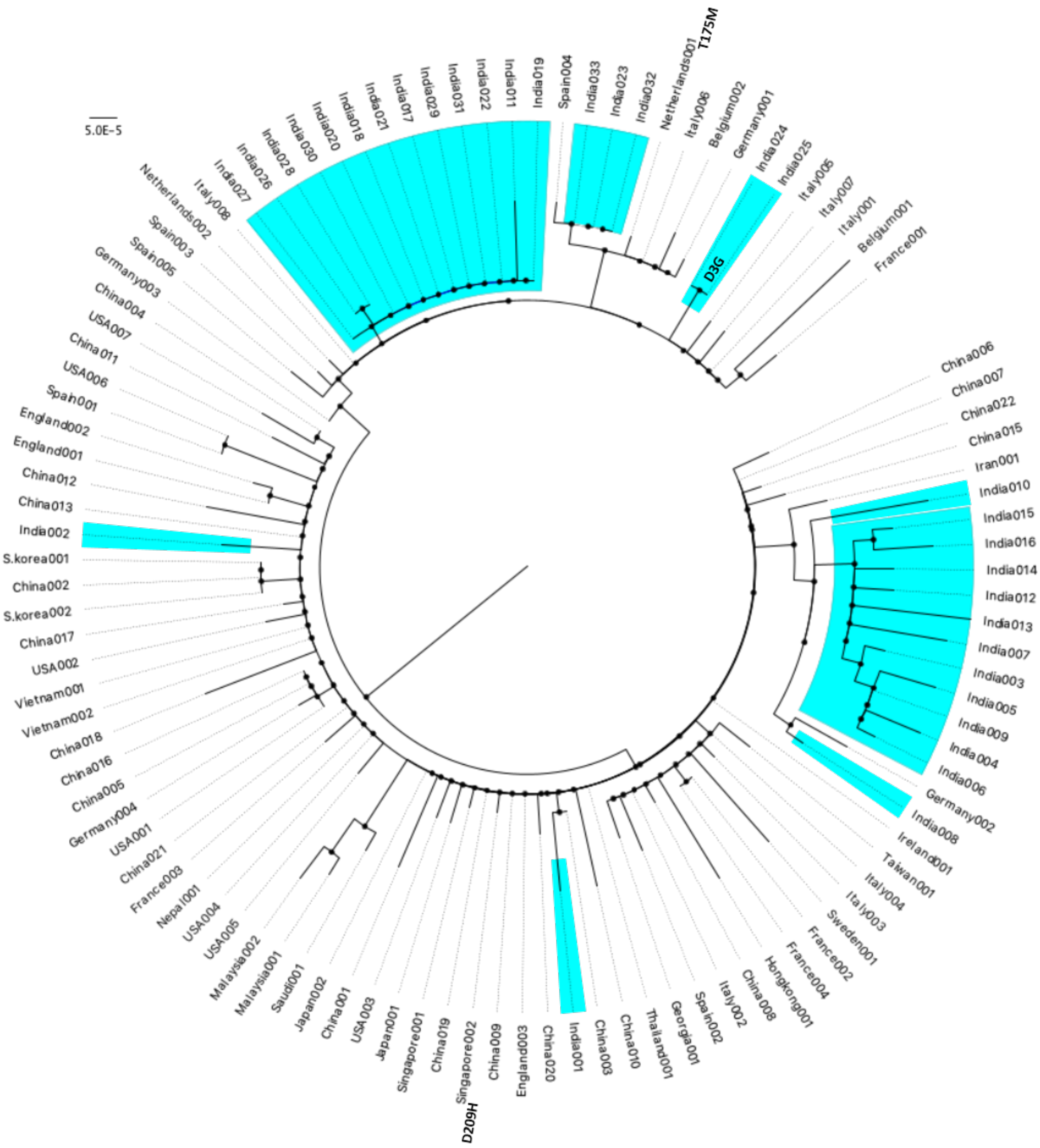


**geneM**

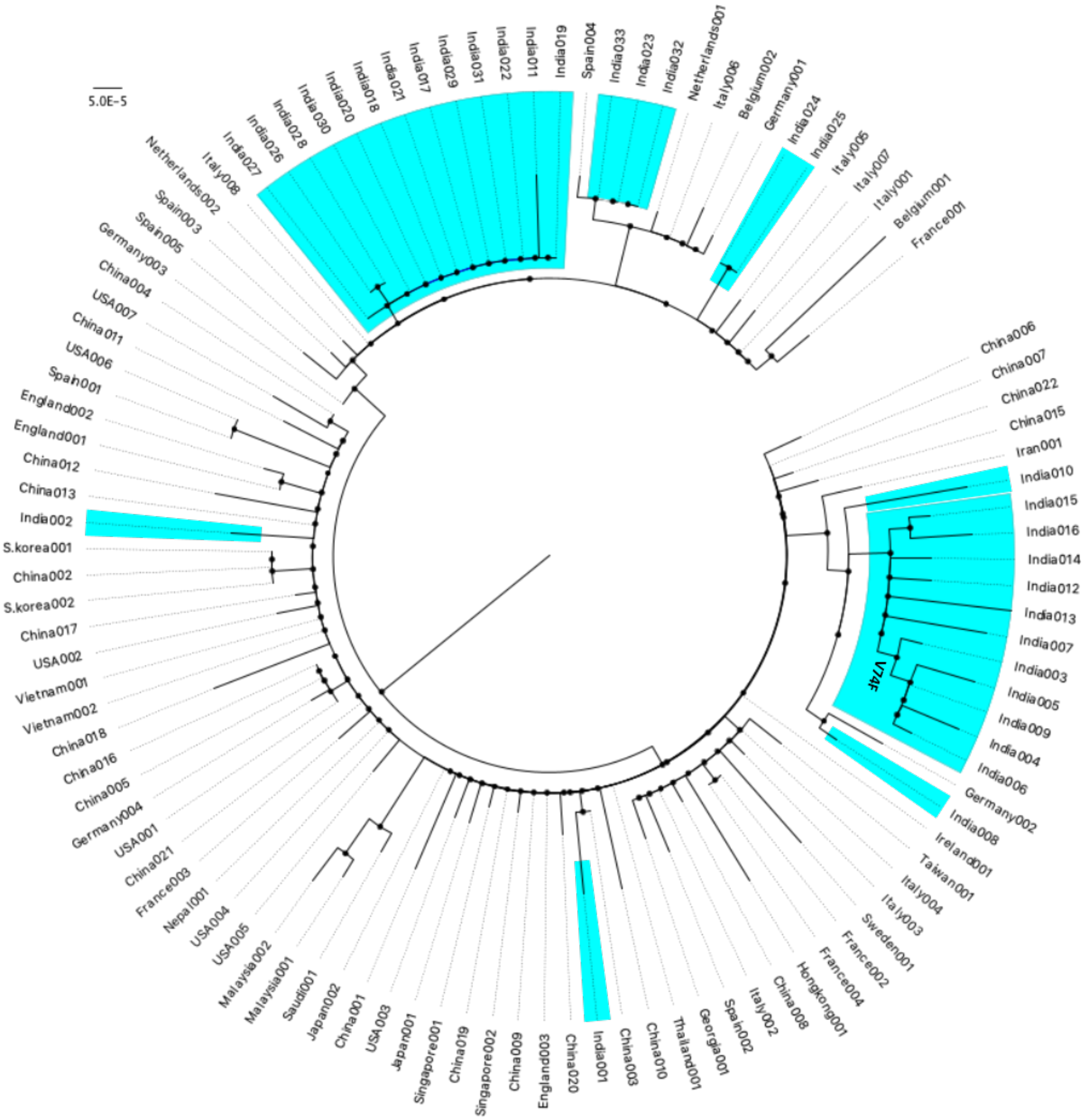
5.0E-5
Netherlands002
Italy008
India027
India026
India028
India030
India020
India018
India021
India017
India029
India031
India022
India011
India019
Spain004
India033
India023
India032
Netherlands001
Italy006
Belgium002
Germany001
India024
India025
Italy005
Italy007
Italy001
Belgium001
France001
China006
China007
China022
China015
Iran001
India010
India015
India016
India014
India012
India013
India007
India003
India005
India009
India004
India006
Germany002
India008
Ireland001
Taiwan001
Italy004
Italy003
Sweden001
France002
France004
Hongkong001
China008
Italy002
Spain002
Georgia001
Thailand001
China010
China003
India001
China020
England003
China009
Singapore002
China019
Singapore001
Japan001
USA003
China001
Japan002
Saudi001
Malaysia001
Malaysia002
USA005
USA004
Nepal001
France003
China021
USA001
Germany004
China005
China016
China018
Vietnam002
Vietnam001
USA002
China017
S.korea002
China002
S.korea001
India002
China013
China012
England001
England002
Spain001
USA006
China011
USA007
China004
Germany003
Spain005
Spain003
V74F

**ORF7a**

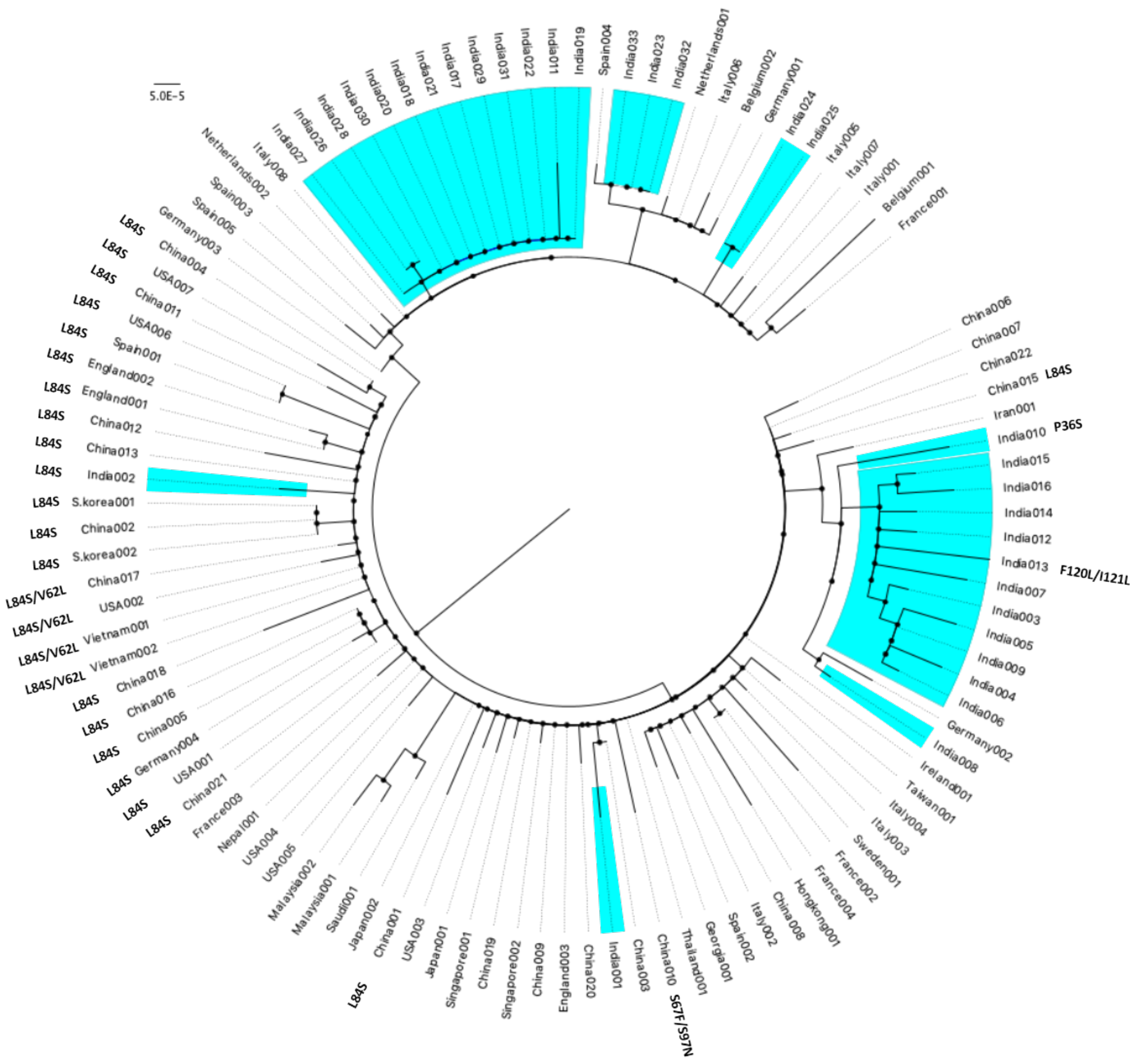

**ORF8**

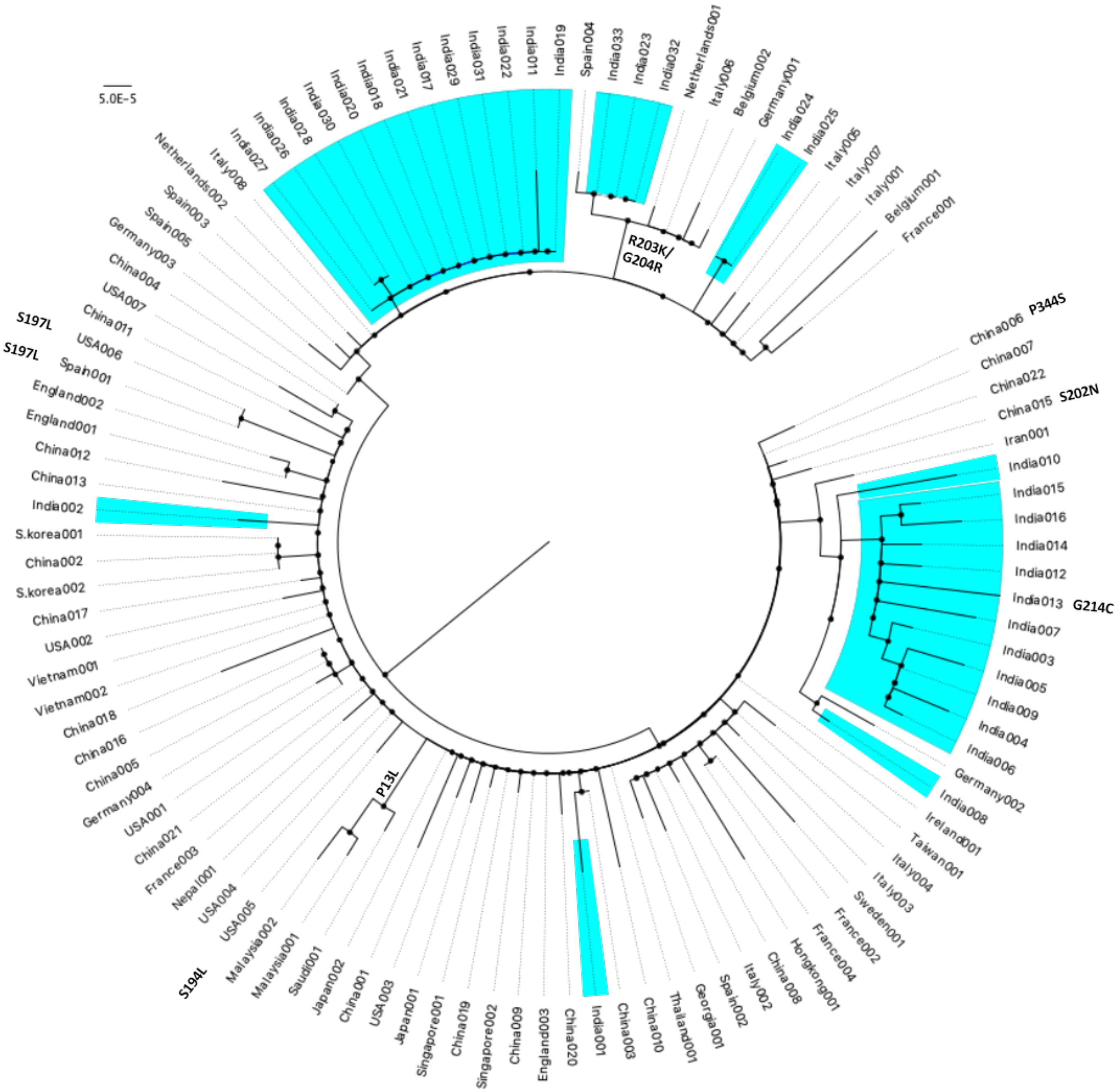

5.0E-5
Italy008
India027
India026
India028
India030
India020
India018
India021
India017
India029
India031
India022
India011
India019
Spain004
India033
India023
India032
Netherlands001
Italy006
Belgium002
Germany001
India024
India025
Italy005
Italy007
Italy001
Belgium001
France001
R203K/
G204R
China006
P344S
China007
China022
China015
S202N
Iran001
India010
India015
India016
India014
India012
India013
G214C
India007
India003
India005
India009
India004
India006
Germany002
India008
Ireland001
Taiwan001
Italy004
Italy003
Sweden001
France002
France004
Hongkong001
China008
Italy002
Spain002
Georgia001
Thailand001
China010
China003
India001
China020
England003
China009
Singapore002
China019
Singapore001
Japan001
USA003
China001
Japan002
Saudi001
Malaysia001
Malaysia002
S194L
USA005
USA004
Nepal001
France003
China021
USA001
Germany004
China005
China016
China018
Vietnam002
Vietnam001
USA002
China017
S.korea002
China002
S.korea001
India002
China013
China012
England001
England002
Spain001
USA006
S197L
China011
S197L
USA007
China004
Germany003
Spain005
Spain003
Netherlands002
P13L


**geneN**

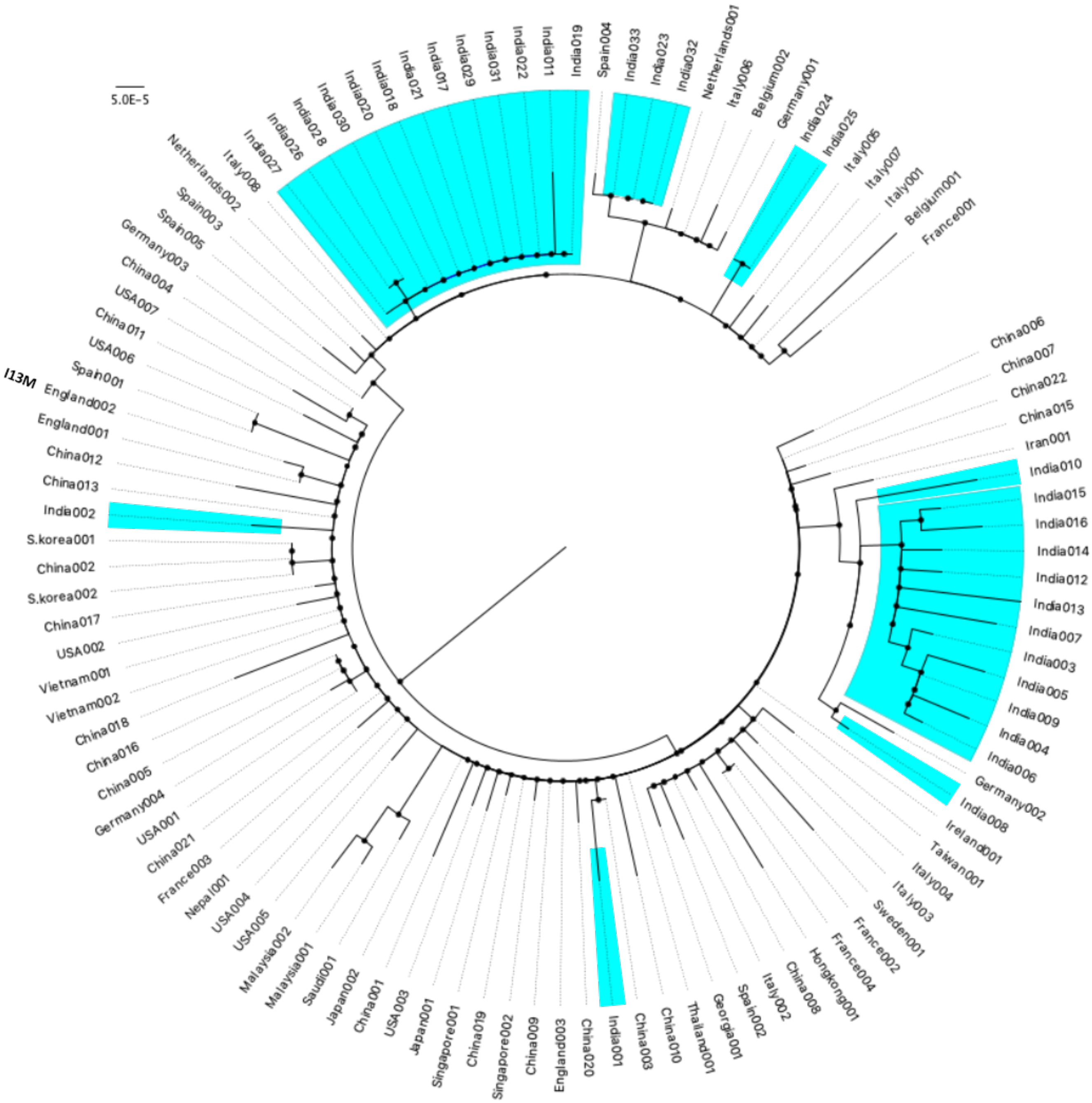
5.0E-5
I13M
Spain004
India033
India023
India032
Netherlands001
Italy06
Belgium002
Germany001
India024
India025
Italy006
Italy007
Italy001
Belgium001
France001
China006
China007
China022
China015
Iran001
India010
India015
India016
India014
India012
India013
India007
India003
India005
India009
India004
India006
Germany002
India008
Ireland001
Taiwan001
Italy004
Italy003
Sweden001
France002
France004
Hongkong001
China008
Italy002
Spain002
Georgia001
Thailand001
China010
China003
India001
China020
England003
China009
Singapore002
China019
Singapore001
Japan001
USA003
China001
Japan002
Saudi001
Malaysia001
Malaysia002
USA005
USA004
Nepal001
France003
China021
USA001
Germany004
China005
China016
China018
Vietnam002
Vietnam001
USA002
China017
S.korea002
China002
S.korea001
India002
China013
China012
England001
England002
Spain001
USA006
China011
USA007
China004
Germany003
Spain005
Spain003
Netherlands002
Italy08
India027
India026
India028
India030
India020
India018
India021
India017
India029
India031
India022
India011
India019

**ORF10**
