## Supplementary Figure 3 for "Emergence of multiple variants of SARS-CoV-2 with signature structural changes"

**Supplementary Figure 3: Secondary structure prediction for spike glycoprotein S**

```
             EEEEEEEECCTCCEEEEEEEEEEEEEEEEEEEEEEHHEEEEHHTHHEHHEHETEEHHHCC
Wuhan        CVADYSVLYNSASFSTFKCYGVSPTKLNDLCFTNVYADSFVIRGDEVRQIAPGQTGKIAD 420
India        CVADYSVLYNSASFSTFKCYGVSPTKLNDLCFTNVYADSFVIRGDEVIQIAPGQTGKIAD 420
             *********************************************** ************
             EEEEEEEECCTCCEEEEEEEEEEEEEEEEEEEEEEHHEEEEHHTHHEEHEHETEEHHHCC

             EEEHHETEEEEEEEECCTTCCCTCCCTCEEEEEEEHHTTHHHHHHTHHHEEEEHCCCCCC
Wuhan        YNYKLPDDFTGCVIAWNSNNLDSKVGGNYNYLYRLFRKSNLKPFERDISTEIYQAGSTPC 480
India        YNYKLPDDFTGCVIAWNFNNLDSKVGGNYNYLYRLFRKSNLKPFERDISTEIYQAGSTPC 480
             ***************** ******************************************
             EEEHHETEEEEEEEEEEECCCCTCCCTCEEEEEEEHHTTHHHHHHTHHHEEEEHCCCCCC

             CEEEEEEEEEEEEEEEEEEEEEHHHHHEEETEEEEEETEEEEEETEEEEEHHHHHCCTCC
Wuhan        GTNTSNQVAVLYQDVNCTEVPVAIHADQLTPTWRVYSTGSNVFQTRAGCLIGAEHVNNSY 660
India        GTNTSNQVAVLYQGVNCTEVPIAIHADQLTPTWRVYSTGSNVFQTRAGCLIGAEHVNNSY 660
             *************.*******:**************************************
             CEEEEEEEEEEEEEEEEEEEEEHHHHHEEETEEEEEETEEEEEETEEEEEHHHHHCCTCC

             HHEEEEEEEEEEEEEHHHHHHEEHTEHHTHHHHHHTHHHTHHHHHHHEEEEEHHHHHHCC
Wuhan        QMAYRFNGIGVTQNVLYENQKLIANQFNSAIGKIQDSLSSTASALGKLQDVVNQNAQALN 960
India        QMAYRFNGIGVTQNVLYENQKLIANQFNSVIGKIQDSLSSTASALGKLQDVVNQNAQALN 960
             *****************************.******************************
             HHEEEEEEEEEEEEEHHHHHHEEHTEEEEEEEHHHTHHHTHHHHHHHEEEEEHHHHHHCC
```

**C -- Coil**
**E -- Sheet**
**H -- Helix**
**T -- Turn**
