## Supplementary Figure 4 for "Emergence of multiple variants of SARS-CoV-2 with signature structural changes"

Supplementary Figure 4: Secondary structure prediction for RdRp protein

```
       EEEEEEEEEHHEEEEEEEEEEEEETTEEEEEEEEEEEEEEEEEEETEEEHHHEEEEHEHT
Wuhan  VNCLDDRCILHCANFNVLFSTVFPPTSFGPLVRKIFVDGVPFVVSTGYHFRELGVVHNQD   300
Indian VNCLDDRCILHCANFNVLFSTVFPLTSFGPLVRKIFVDGVPFVVSTGYHFRELGVVHNQD   300
       ************************ ***********************************
       EEEEEEEEEHHEEEEEEEEEEEEEEEEEEEEEEEEEEEEEEEEEEEETEEEHHHEEEEHEHT

       CCCCCTHHHHHHEEEHHHHHHHHHHHTHHHHHEEEEEEHHHEEEEEEEEEEHCTTCCCCT
Wuhan  VNLHSSRLSFKELLVYAADPAMHAASGNLLLDKRTTCFSVAALTNNVAFQTVKPGNFNKD   360
Indian VNLHSSRLSFKELLVYAADPAMHAASGNLLLDKRTTCFSVAALTNNVVFQTVKPGNFNKD   360
       ***********************************************.************
       CCCCCTHHHHHHEEEHHHHHHHHHHHTHHHHHEEEEEEHHHEEEEEEEEEEHCTTCCCCT

       CCCEHEEEEEHEEEEHHHHEEEEHHHHCCCCCCCCTHHHHHEEEEEEEEEHHHHHHEECC
Wuhan  GAGCFVDDIVKTDGTLMIERFVSLAIDAYPLTKHPNQEYADVFHLYLQYIRKLHDELMLT   840
Indian GAGCFVDDIVKTDGTLMIERFVSLAIDAYPLTKHPSQEYADVFHLYLQYIRKLHDELMLT   840
       ***********************************.************************
       CCCEHEEEEEHEEEEHHHHEEEEHHHHCCCCCCCCTHHHHHEEEEEEEEEHHHHHHEECC
```
