## Supplementary Table 1 for "Emergence of multiple variants of SARS-CoV-2 with signature structural changes"

Supplementary Table 1: List of all 109 complete genomes considered in this study.

| Sequence ID | Region | Accession ID | Source | Collection Date |
|---|---|---|---|---|
| Belgium001 | Europe/Belgium/Leuven | EPI_ISL_415159 | www.gisaid.org | 2020-02-29 |
| Belgium002 | Europe/Belgium/Brussels | EPI_ISL_415158 | www.gisaid.org | 2020-03-01 |
| China001 | Asia/China/Anhui | EPI_ISL_413485 | www.gisaid.org | 2020-01-24 |
| China002 | Asia/China/Beijing | EPI_ISL_413519 | www.gisaid.org | 2020-01-28 |
| China003 | Asia/China/Guangdong | EPI_ISL_406535 | www.gisaid.org | 2020-01-22 |
| China004 | Asia/China/Fujian | EPI_ISL_411060 | www.gisaid.org | 2020-01-21 |
| China005 | Asia/China/Guandong | EPI_ISL_403932 | www.gisaid.org | 2020-01-14 |
| China006 | Asia/China/Guangdong | EPI_ISL_406533 | www.gisaid.org | 2020-01-22 |
| China007 | Asia/China/Zhejiang | EPI_ISL_407313 | www.gisaid.org | 2020-01-19 |
| China008 | Asia/China/Hangzhou | EPI_ISL_415709 | www.gisaid.org | 2020-01-25 |
| China009 | Asia/China/Anhui | EPI_ISL_412026 | www.gisaid.org | 2020-02-23 |
| China010 | Asia/China/Hubei | EPI_ISL_412459 | www.gisaid.org | 2020-01-08 |
| China011 | Asia/China/Jiangxi | EPI_ISL_421241 | www.gisaid.org | 2020-01-30 |
| China012 | Asia/China/Jiangxi | EPI_ISL_421245 | www.gisaid.org | 2020-01-25 |
| China013 | Asia/China/Jiangxi | EPI_ISL_421246 | www.gisaid.org | 2020-02-26 |
| China015 | Asia/China/Shanghai | EPI_ISL_416316 | www.gisaid.org | 2020-01-25 |
| China016 | Asia/China/Guangdong | EPI_ISL_406030 | www.gisaid.org | 2020-01-10 |
| China017 | Asia/China/Sichuan | EPI_ISL_408484 | www.gisaid.org | 2020-01-15 |
| China018 | Asia/China/Hubei | EPI_ISL_412983 | www.gisaid.org | 2020-02-08 |
| **China019*** | **Asia/China/Hubei/Wuhan** | **NC_045512.2** | NCBI | 2019-12 |
| China020 | Asia/China/Hubei | EPI_ISL_402130 | www.gisaid.org | 2019-12-30 |
| China021 | Asia/China/Yunnan | EPI_ISL_408480 | www.gisaid.org | 2020-01-17 |
| China022 | Asia/China/Zhejiang | EPI_ISL_404227 | www.gisaid.org | 2020-01-16 |
| England001 | Europe/England | EPI_ISL_407071 | www.gisaid.org | 2020-01-29 |
| England002 | Europe/England | EPI_ISL_407073 | www.gisaid.org | 2020-01-29 |
| England003 | Europe/England | EPI_ISL_414040 | www.gisaid.org | 2020-02-05 |
| England004 | Europe/England/Yorkshire | EPI_ISL_414500 | www.gisaid.org | 2020-03-04 |
| France001 | Europe/France/Hauts-de-France | EPI_ISL_414638 | www.gisaid.org | 2020-03-04 |
| France002 | Europe/France/Ile-de-France | EPI_ISL_406596 | www.gisaid.org | 2020-01-23 |
| France003 | Europe/France/Ile-de-France | EPI_ISL_411218 | www.gisaid.org | 2020-02-02 |
| France004 | Europe/France/Ile-de-France | EPI_ISL_411220 | www.gisaid.org | 2020-01-28 |
| Georgia001 | Europe/Georgia/Tbilisi | EPI_ISL_415643 | www.gisaid.org | 2020-03-10 |
| Germany001 | Europe/Germany/Baden-Wuerttemberg | EPI_ISL_412912 | www.gisaid.org | 2020-02-25 |
| Germany002 | Europe/Germany/Munich | EPI_ISL_414520 | www.gisaid.org | 2020-03-02 |
| Germany003 | Europe/Germany/Munich | EPI_ISL_406862 | www.gisaid.org | 2020-01-28 |
| Germany004 | Europe/Germany/Munich | EPI_ISL_414521 | www.gisaid.org | 2020-03-02 |
| Hongkong001 | Asia/Hongkong | EPI_ISL_412029 | www.gisaid.org | 2020-01-30 |
| India001 | Asia/India/Kerala | EPI_ISL_413522 | www.gisaid.org | 2020-01-27 |
| India002 | Asia/India/Kerala | EPI_ISL_413523 | www.gisaid.org | 2020-01-31 |
| India003 | Asia/India | EPI_ISL_424361 | www.gisaid.org | 2020-03-10 |
| India004 | Asia/India | EPI_ISL_421662 | www.gisaid.org | 2020-03-10 |
| India005 | Asia/India | EPI_ISL_421663 | www.gisaid.org | 2020-03-10 |
| India006 | Asia/India | EPI_ISL_421664 | www.gisaid.org | 2020-03-10 |

| | | | | |
|---|---|---|---|---|
| India007 | Asia/India | EPI_ISL_421665 | www.gisaid.org | 2020-03-10 |
| India008 | Asia/India | EPI_ISL_421666 | www.gisaid.org | 2020-03-10 |
| India009 | Asia/India | EPI_ISL_421667 | www.gisaid.org | 2020-03-10 |
| India010 | Asia/India | EPI_ISL_421668 | www.gisaid.org | 2020-03-10 |
| India011 | Asia/India | EPI_ISL_424362 | www.gisaid.org | 2020-03-10 |
| India012 | Asia/India | EPI_ISL_421669 | www.gisaid.org | 2020-03-12 |
| India013 | Asia/India | EPI_ISL_421670 | www.gisaid.org | 2020-03-12 |
| India014 | Asia/India | EPI_ISL_421671 | www.gisaid.org | 2020-03-12 |
| India015 | Asia/India | EPI_ISL_421672 | www.gisaid.org | 2020-03-12 |
| India016 | Asia/India | EPI_ISL_424363 | www.gisaid.org | 2020-03-12 |
| India017 | Asia/India | EPI_ISL_420544 | www.gisaid.org | 2020 |
| India018 | Asia/India | EPI_ISL_420546 | www.gisaid.org | 2020 |
| India019 | Asia/India | EPI_ISL_420548 | www.gisaid.org | 2020 |
| India020 | Asia/India | EPI_ISL_420550 | www.gisaid.org | 2020 |
| India021 | Asia/India | EPI_ISL_420552 | www.gisaid.org | 2020 |
| India022 | Asia/India | EPI_ISL_420554 | www.gisaid.org | 2020 |
| India023 | Asia/India | EPI_ISL_420556 | www.gisaid.org | 2020 |
| India024 | Asia/India | EPI_ISL_424364 | www.gisaid.org | 2020-03-17 |
| India025 | Asia/India | EPI_ISL_424365 | www.gisaid.org | 2020-03-17 |
| India026 | Asia/India | EPI_ISL_420543 | www.gisaid.org | 2020-03-03 |
| India027 | Asia/India | EPI_ISL_420545 | www.gisaid.org | 2020-03-03 |
| India028 | Asia/India | EPI_ISL_420547 | www.gisaid.org | 2020-03-03 |
| India029 | Asia/India | EPI_ISL_420549 | www.gisaid.org | 2020-03-03 |
| India030 | Asia/India | EPI_ISL_420551 | www.gisaid.org | 2020-03-03 |
| India031 | Asia/India | EPI_ISL_420553 | www.gisaid.org | 2020-03-03 |
| India032 | Asia/India | EPI_ISL_426179 | www.gisaid.org | 2020-03-02 |
| India033 | Asia/India | EPI_ISL_420555 | www.gisaid.org | 2020-03-03 |
| Iran001 | Asia/Iran | MT320891.2 | NCBI | 2020-03-09 |
| Ireland001 | Europe/Ireland/Cork | EPI_ISL_414487 | www.gisaid.org | 2020-03-04 |
| Italy001 | Europe/Italy/Lombardy | EPI_ISL_412973 | www.gisaid.org | 2020-02-20 |
| Italy002 | Europe/Italy/Rome | EPI_ISL_410546 | www.gisaid.org | 2020-01-31 |
| Italy003 | Europe/Italy/Rome | EPI_ISL_410545 | www.gisaid.org | 2020-01-29 |
| Italy004 | Europe/Italy/Rome | EPI_ISL_412974 | www.gisaid.org | 2020-01-29 |
| Italy005 | Europe/Italy/Lombardy | EPI_ISL_413489 | www.gisaid.org | 2020-03-03 |
| Italy006 | Europe/Italy/Rome | EPI_ISL_417922 | www.gisaid.org | 2020-02-28 |
| Italy007 | Europe/Italy/Abruzzo | EPI_ISL_418255 | www.gisaid.org | 2020-03-14 |
| Italy008 | Europe/Italy/Lombardy | EPI_ISL_417447 | www.gisaid.org | 2020-02-24 |
| Japan001 | Asia/Japan/Aichi | EPI_ISL_407084 | www.gisaid.org | 2020-01-25 |
| Japan002 | Asia/Japan/Nara | EPI_ISL_410531 | www.gisaid.org | 2020-01-25 |
| Malaysia001 | Asia/Malaysia/Kuala Lumpur | EPI_ISL_417918 | www.gisaid.org | 2020-03-18 |
| Malaysia002 | Asia/Malaysia/Kuala Lumpur | EPI_ISL_417917 | www.gisaid.org | 2020-03-20 |
| Nepal001 | Asia/Nepal/Kathmandu | EPI_ISL_410301 | www.gisaid.org | 2020-01-13 |
| Netherlands001 | Europe/Netherlands/Gelderland | EPI_ISL_414423 | www.gisaid.org | 2020-03-02 |
| Netherlands002 | Europe/Netherlands/Noord | EPI_ISL_414428 | www.gisaid.org | 2020-03-02 |
| S.korea001 | Asia/South Korea/Gyeonggi-do | EPI_ISL_407193 | www.gisaid.org | 2020-01-25 |
| S.korea002 | Asia/South Korea | EPI_ISL_413018 | www.gisaid.org | 2020-02-06 |
| Saudi001 | Asia/Saudi/Riyadh | EPI_ISL_416432 | www.gisaid.org | 2020-03-07 |

| | | | | |
|---|---|---|---|---|
| Singapore001 | Asia/Singapore | EPI_ISL_406973 | www.gisaid.org | 2020-01-23 |
| Singapore002 | Asia/Singapore | EPI_ISL_407987 | www.gisaid.org | 2020-01-25 |
| Spain001 | Europe/Spain/Comunitat | EPI_ISL_414598 | www.gisaid.org | 2020-03-05 |
| Spain002 | Europe/Spain/Andalusia | EPI_ISL_418243 | www.gisaid.org | 2020-02-28 |
| Spain003 | Europe/Spain/Castilla | EPI_ISL_418247 | www.gisaid.org | 2020-02-26 |
| Spain004 | Europe/Spain/Catalonia | EPI_ISL_418250 | www.gisaid.org | 2020 |
| Spain005 | Europe/Spain/Madrid | EPI_ISL_418251 | www.gisaid.org | 2020-02-25 |
| Sweden001 | Europe/Sweden | EPI_ISL_411951 | www.gisaid.org | 2020-02-07 |
| Taiwan001 | Asia/Taiwan/Kaohsiung | EPI_ISL_406031 | www.gisaid.org | 2020-01-23 |
| Thailand001 | Asia/Thailand/Nonthaburi | EPI_ISL_403962 | www.gisaid.org | 2020-01-08 |
| USA001 | North America/USA/Arizona | EPI_ISL_406223 | www.gisaid.org | 2020-01-22 |
| USA002 | North America/USA/California | EPI_ISL_406034 | www.gisaid.org | 2020-01-23 |
| USA003 | North America/USA | EPI_ISL_413606 | www.gisaid.org | 2020-02-17 |
| USA004 | North America/USA/Illinois | EPI_ISL_404253 | www.gisaid.org | 2020-01-21 |
| USA005 | North America/USA/Massachusetts | EPI_ISL_409067 | www.gisaid.org | 2020-01-29 |
| USA006 | North America/USA/New York | EPI_ISL_415151 | www.gisaid.org | 2020-03-04 |
| USA007 | North America/USA/Washington | EPI_ISL_415622 | www.gisaid.org | 2020-03-09 |
| Vietnam001 | Asia/Vietnam/Thanhhoa | EPI_ISL_416427 | www.gisaid.org | 2020-01-24 |
| Vietnam002 | Asia/Vietnam/Thanhhoa | EPI_ISL_408668 | www.gisaid.org | 2020-01-24 |

*** reference genome**
